# Dynamic protein quantitation (DyProQ) of procollagen-I by CRISPR-Cas9 NanoLuciferase tagging

**DOI:** 10.1101/2020.05.17.099119

**Authors:** Ben C. Calverley, Karl E. Kadler, Adam Pickard

## Abstract

The ability to quantitate a protein of interest temporally and spatially at subcellular resolution in living cells would generate new opportunities for research and drug discovery but remains a major technical challenge. Here, we describe dynamic protein quantitation (DyProQ) which is effective across microscopy and multiwell platforms. Using collagen as a test protein, CRISPR-Cas9-mediated introduction of nluc (encoding NanoLuciferase, NLuc) into the *Col1a2* locus enabled simplification and miniaturisation of procollagen-I (PC-I) quantitation. We robustly assessed extracellular, intracellular, and subcellular PC-I levels, by correlating to known concentrations of recombinant NLuc in the presence of substrate. Loss of collagen causes tissue degeneration whereas excess collagen results in fibrosis (often with poor-outcome) and is evident in aggressive cancers; however, treatment options are extremely limited. Using collagen-DyProQ, we screened a library of 1,971 FDA-approved compounds and identified 10 candidates for repurposing in the treatment of fibrotic and 7 for degenerative diseases.

---

Quantitation of DNA and RNA is routine in research and diagnostic laboratories and makes use of base pair hybridization to ensure specificity and identification. Similar approaches are not available for proteins. Methods such as ELISA immunoassays and western blotting are widely used to estimate levels of proteins, but spatial resolution is lost, and they are unsuitable for live cell studies where dynamic readouts are required. In this regard, the use of fluorescent proteins and chemical tags has revolutionized cell biology, but quantitation through fluorescence is not without technical difficulties associated with quenching, sometimes extensive wash-out, and the influence of local environment on the fluorescence signal. Low fluorescence signals can be overcome with the use of strong exogenous promoters, but these disrupt the endogenous behaviour of the protein under study.

Bioluminescence produced when luciferase hydrolyses luciferin-based substrates offers a practical alternative to using fluorescent tags. When tagged to a protein of interest, luciferase emits visible light in the presence of a suitable substrate. Hall *et al.* used a small luciferase subunit from the deep-sea shrimp *Oplophorus gracilirostris* to produce NanoLuciferase (NLuc), which produces more photons than either firefly or Renilla luciferases when used in combination with a novel imidazopyrazinone substrate, furimazine^1^. In our study we used CRISPR-Cas9 to fuse NLuc to the N-terminus of procollagen-I (PC-I), which is the precursor of collagen-I and the most abundant protein in vertebrates^2^. Collagen-I is a triple helical protein^3^ that occurs in the extracellular matrix as elongated fibrils that are established during development^4^ and remain throughout adulthood without turnover^5^ in the presence of a sacrificial pool of collagen that is under circadian control^6^. Although the scaffolding function of collagen I is essential for tissue integrity, excess collagen causes tissue damage in fibrosis (scarring) and is associated with aggressive cancers^7,8^ and 45% of deaths^9^. Thus, collagen is of broad clinical importance, from regenerative medicine in which elevating collagen synthesis is needed to build tissue, to fibrosis in which inhibiting collagen synthesis is required to stop loss of tissue function. However, the identification of drugs to either increase or decrease collagen levels is hampered by the lack of suitable technologies for measuring collagen levels in cell culture. Collagen-I contains ~ 13.6% hydroxyproline^10^, and assay of hydroxyproline has become the gold standard for quantifying tissue collagen. However, hydroxyproline also occurs in the 27 other collagens^11^, non-collagenous triple helical proteins (reviewed by ^12^), and elastin^13^, which is difficult to take into account when using hydroxyproline to estimate levels of collagen-I. Moreover, the assay is destructive and unsuitable for time-resolved studies of collagen synthesis in single cells. Proteomics^6^, western blotting, and the use of fluorescent tags (e.g. green fluorescence protein, GFP) are either destructive or require the use of overexpression promoters to provide good signal/noise ratios. Furthermore, these approaches cannot quantify the rapid synthesis and secretion of collagen, which pulse-chase approaches (using ^3^H- and ^14^C-biosynthetic labelling) have shown can occur within mins^14^. In our study, we show that the light produced by NLuc is sufficiently bright to obtain dynamic quantitative information on the number of endogenous PC-I molecules trafficking through living cells and could be ported to a 96-well format with which we screened a collection of FDA-approved compounds to identify compounds that are effective regulators of collagen synthesis and secretion.

## Results

### CRISPR-Cas9 editing of *Col1a2*

We designed a multifunctional tag to aid the identification and clonal selection of *nluc::Col1a2* fibroblasts. The tag retains the ER-targeting signal recognition sequence (SP) of *Col1a2*, GFP11 sequences from GFP for use in split-GFP approaches, 6 histidine residues for PC-I capture, and NLuc (Fig. 1A). Selection of edited cells was achieved using fibroblasts expressing the GFP barrel (Fig. 1B). Confirmation of CRISPR editing was confirmed by PCR from genomic DNA (Fig. 1C) and quantitation and sequencing of RNA transcripts across the junctions of *nluc* and *Col1a2* (Fig. 1D). Sequencing of PCR products confirmed introduction of *nluc* in-frame with *Col1a2* (Fig. 1E). Secretion of NLuc-PC-I was confirmed by His-trap capture of the protein from the medium of edited cells (Supplementary Fig. 1). A peptide spanning the junction of NLuc and proα2(I) was identified by LC-MS/MS (Fig. 1F). Incorporation of NLuc into the heterotrimer of PC-I was also confirmed in high molecular weight complexes, where association with proα1(I) was identified (Supplementary Fig. 1). Under reduced conditions, NLuc-PC-I was identified by in-gel detection of NLuc activity at approximately 120 kDa (Fig. 1G). The culture medium from *nluc::Col1a2* fibroblasts was passed over a Ni^2+^ chelating column and bound proteins were eluted with imidazole. The fractions were separated by SDS-PAGE, the gel was stained with Coomassie blue and protein bands subjected to LC-MS/MS for protein identification (Supplementary Fig. 1). The results showed the presence of intact (His)6-NLuc-PC-I, (His)6-NLuc-pCcollagen-I, and collagen-I. The presence of free (His)6-NLuc showed that N-proteinase was capable of cleaving (His)6-NLuc-PC-I, which is a good indicator that NLuc-PC-I secreted by *nluc::Col1a2* fibroblasts was triple helical^15^.

**Figure 1:**
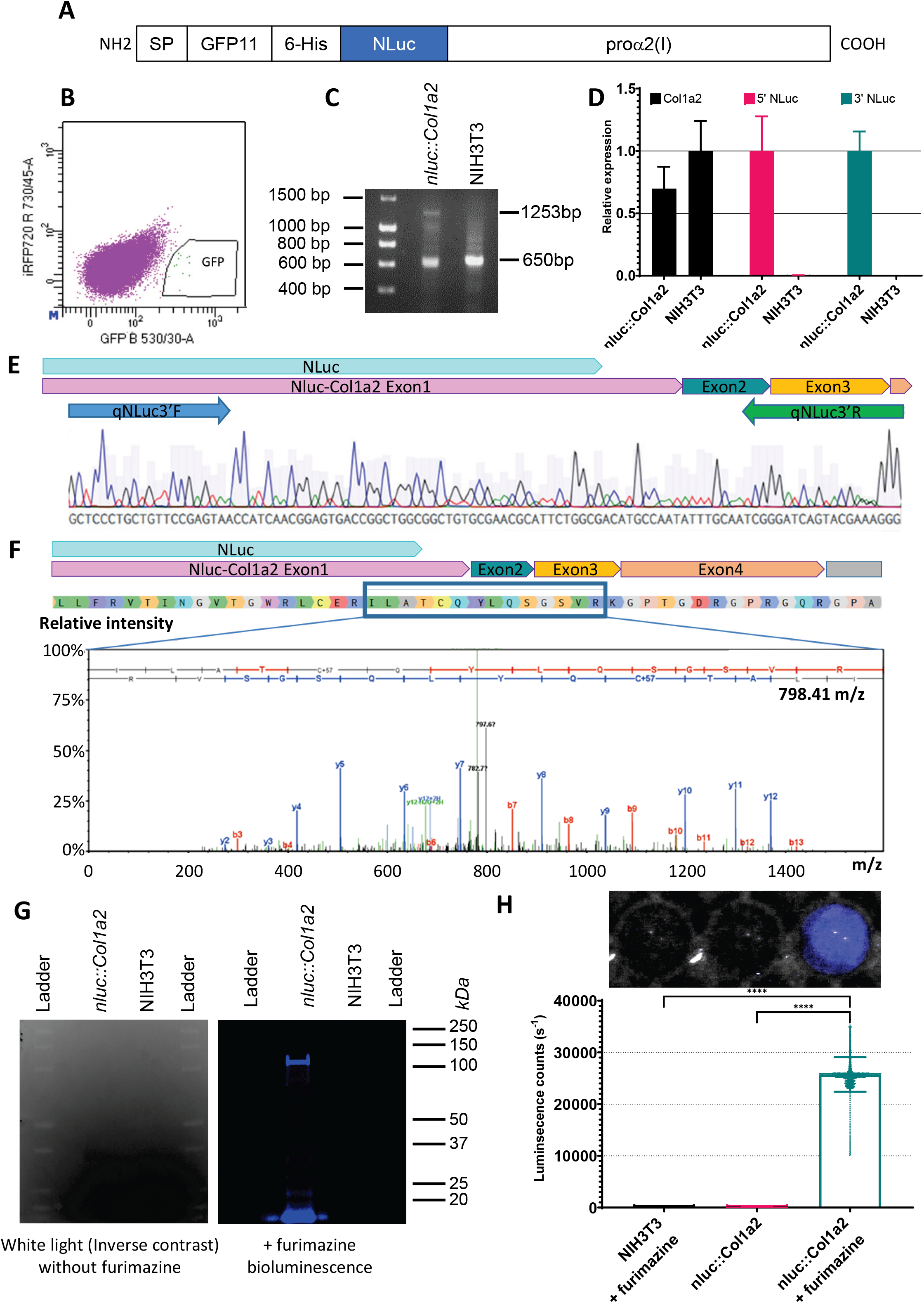
Quantitative CRISPR/Cas9 editing strategy. **A)** Tagging strategy to allow quantitation of PC-I in NIH3T3 fibroblasts with introduction of a multifunctional tag at the N-terminus of proα2(I) to allow detection of edited cells, and the small luciferase, NLuc. **B)** After introduction of the multifunctional tag in exon 1 of *Col1a2* using CRISPR/Cas9, edited cells were sorted based on GFP fluorescence. **C)** PCR validation of edited DNA in cells isolated from B) using primers in Supplementary table 1. **D)** Real-time PCR of total *Col1a2* transcripts (Black) and edited transcripts in NIH3T3 and *nluc::Col1a2* cells. Primers at the 5’ and 3’ ends of the introduced nluc confirmed insertion into the *Col1a2* transcript. Bars show mean ± SD, n=3 independent experiments. **E)** Boxes show exon positions of the edited *Col1a2* RNA transcript, sequencing of PCR products in D) demonstrate insertion of *nluc* in frame with *Col1a2*. **F)** Boxes show exon positions of the edited *Col1a2* RNA transcript and the amino acid sequence of the expected tagged protein. Conditioned medium from *nluc::Col1a2* cells was affinity purified (Supplementary Fig. 1) and subjected to mass spectrometry analysis. A peptide spanning NLuc and protein encoded by Col1a2 exons 1-3 was identified. **G)** In-gel detection of NLuc tagged proα2(I) chain under reducing conditions identified NLuc activity at approximately 140 kDa. **H)** Imaging and quantitation of the light produced by *nluc::Col1a2* and unedited NIH3T3 cells incubated with the NLuc substrate, Furimazine (Nano-Glo). On a 96-well plate, single wells were imaged using a single-lens reflex camera and quantitation of photon counts using a multi-well plate luminometer. Bar shows means ± SD from n=30 replicate measurements. **** represents p=0.0001, paired Student’s t-Test.

To demonstrate the ease of detecting NLuc-PC-I, an SLR camera was used to capture the light produced by a single well of a 96-well plate containing *nluc::Col1a2* cells, following addition of Furimazine (Fig. 1H). As endogenous NLuc activity generated a remarkably bright signal we were required to optimise the plasticware for the assay. Black, white, and clear plates were tested. The results showed that white plates reduced spill over of light between wells whilst maximising emission, and were therefore used in all subsequent 96-well plate reader experiments (Supplementary Fig. 2).

### Quantitation of NLuc-PC-I

As a first experiment, we used the chloramine-T colorimetric method to quantify the amount of hydroxyproline synthesized by *nluc::Col1a2* fibroblasts. In our hands, at least 300,000 cells were required to synthesise sufficient collagen to be detected using this method (Fig. 2A). Next, we compared known numbers of *nluc::Col1a2* cells and known numbers of matched 3T3 cells, and measured hydroxyproline in the cell layer from each set. The results showed that CRISPR-Cas9 editing of the cells did not alter the ability of the cells to synthesise collagen (Fig. 2B). We cultured *nluc::Col1a2* cells, added Furimazine to the cells, and measured the resultant luminescence (Fig. 2C). These experiments demonstrated the high sensitivity of NLuc detection, compared to measurement of hydroxyproline, to detect PC-I synthesis. To be able to quantitate the number of NLuc-PC-I molecules synthesised per cell, we prepared a standard curve of luminescence from recombinant NLuc (rNLuc) in the presence of Furimazine (Fig. 2D). An important consideration was whether we could infer a direct correlation between luminescence produced by rNLuc in a well-mixed solution, and NLuc bound to collagen within cells and subcellular compartments. Direct comparison of lysed and un-lysed cells indicated there is no significant difference in the time taken for luminescence levels to peak following addition of Furimazine (Supplementary Fig. 3). Differences in the absolute level of luminescence were observed; this was explained by differences in the activity of rNLuc in lysis buffer versus DMEM medium. The outcome of these experiments was confidence that we could correlate luminescence levels recorded from known numbers of NLuc molecules to luminescence levels recorded from unknown numbers of NLuc-PC-I molecules in cells or culture medium. By bringing the luminescence and cell number data together, we were able to describe a relationship converting luminescence to numbers of NLuc-PC-I molecules and showed that luminescence was linear over a range of 78 to 4,000,000 cells (Fig. 2E). Of note, the luminescence counts per rNLuc molecule were constant over a wide range of rNLuc molecules without noticeable quenching or amplification (Fig. 2F). Furthermore, a consistent value of 228,000 (median, 3 s.f.) and 225,000 (mean, given by dashed line in Fig. 2E, 3 s.f.) was obtained for numbers of NLuc-PC-I molecules per cell after correlation of luminescence from cells and rNLuc across 5 orders of magnitude.

**Figure 2:**
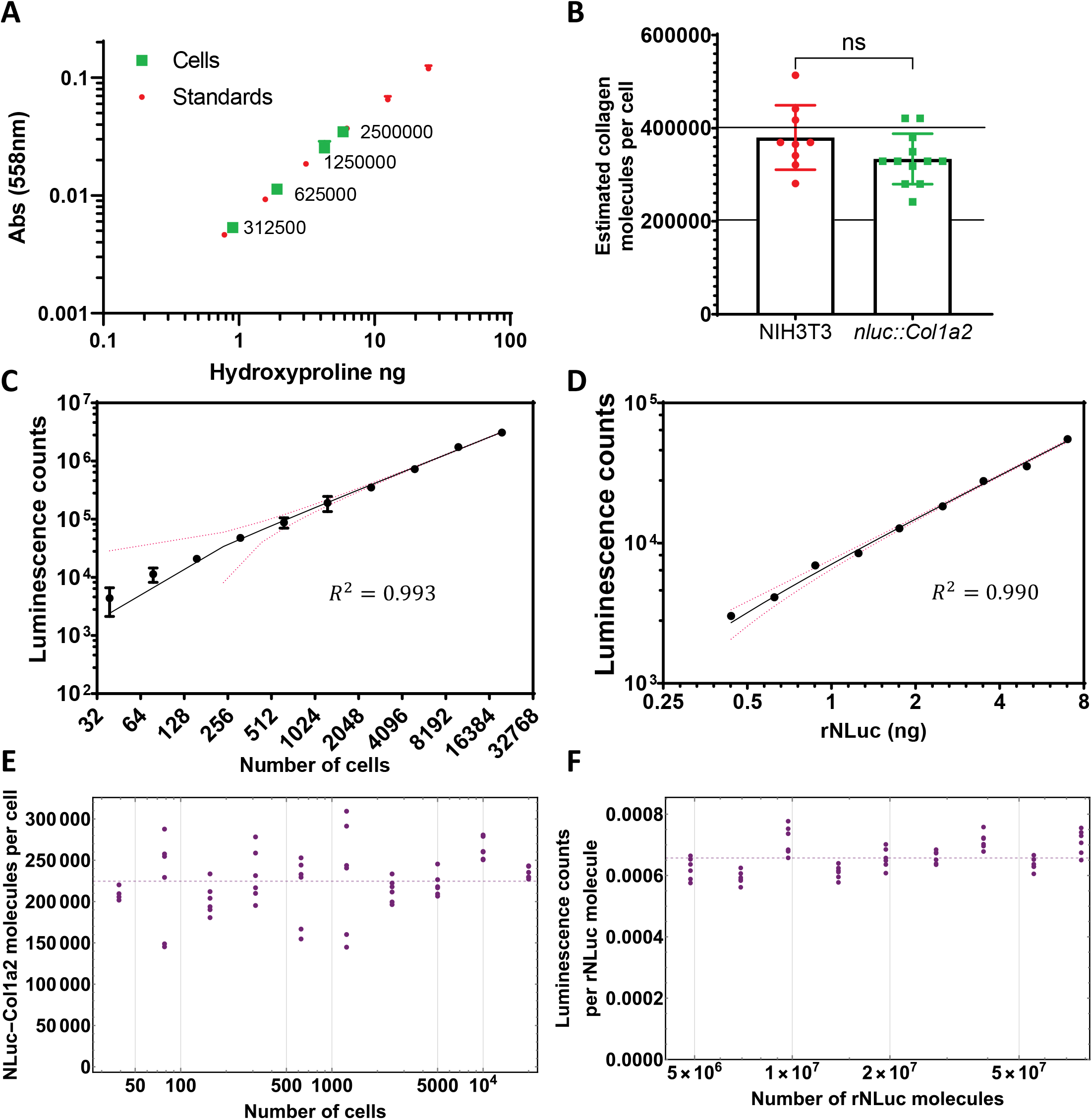
Quantitation of intracellular collagen molecules. **A)** The quantity of hydroxyproline in known numbers of cells were calculated using known concentrations of hydroxyproline. Greater than 300,000 cells were required to ensure accurate comparison to hydroxyproline standards. **B)** Estimated numbers of collagen molecules per cell based on hydroxyproline. n=3 independent experiments each conducted in triplicate, NIH3T3 and n=4 independent experiments each conducted in triplicate. Bars show mean ± SD. **C)** Bioluminescence counts per second of *nluc::Col1a2* cells scales with cell number. n=3 independent assays, each recorded n=4 times **D)** Correlation of known quantities of recombinant NLuc (rNLuc) with luminescence counts. n=3 independent assays, each recorded n=4 times. **E)** Across the range of rNLuc concentrations tested, reaction conditions allowed consistent counts per rNLuc molecule to be measured. **F)** By comparing the bioluminescence from C, and D the number of NLuc-PC-I molecules per cell could be quantified.

### Direct imaging and quantitation of NLuc-PC-I in cells

Next, we wanted to know if we could quantitate numbers of NLuc-PC-I molecules in bioluminescence microscopy images of the cells. This would provide quantitative information on PC-I trafficking and allow us to assess the sensitivity of collagen-DyProQ. By correlating bioluminescence from known amounts of rNLuc (Supplementary Fig. 4A-C), we were able to determine the number of NLuc-PC-I molecules in bioluminescence images. The total luminescence in each cell within the field of view could then be individually calculated and converted to the total number of NLuc-PC-I molecules per cell (Fig. 3). The results showed a mean of 207,000 (3 s.f.) and a median of 229,000 (3 s.f.) NLuc-PC-I molecules per cell (Fig. 3B). These estimates of NLuc-PC-I molecules per cell from bioimages were in strong agreement with the estimates obtained using a plate reader (differing by less than 10% in mean values, and less than 1% in median values). A range of 111,000 to 290,000 (3 s.f.) NLuc-PC-I molecules per cell was observed, representing a 62% variation in cellular collagen levels. We noticed bright luminescence in subcellular vesicles (see for example highlighted region in Fig. 3C). From measurements of photon counts we were able to estimate ~10,800 NLuc-PC-I molecules in the vesicle shown. If we assume the vesicle to be spherical (diameter 4.15 μm (3 s.f.)) then the concentration of NLuc-PC-I within this vesicle is ~ 0.231 mg/mL (3s.f.).

**Figure 3:**
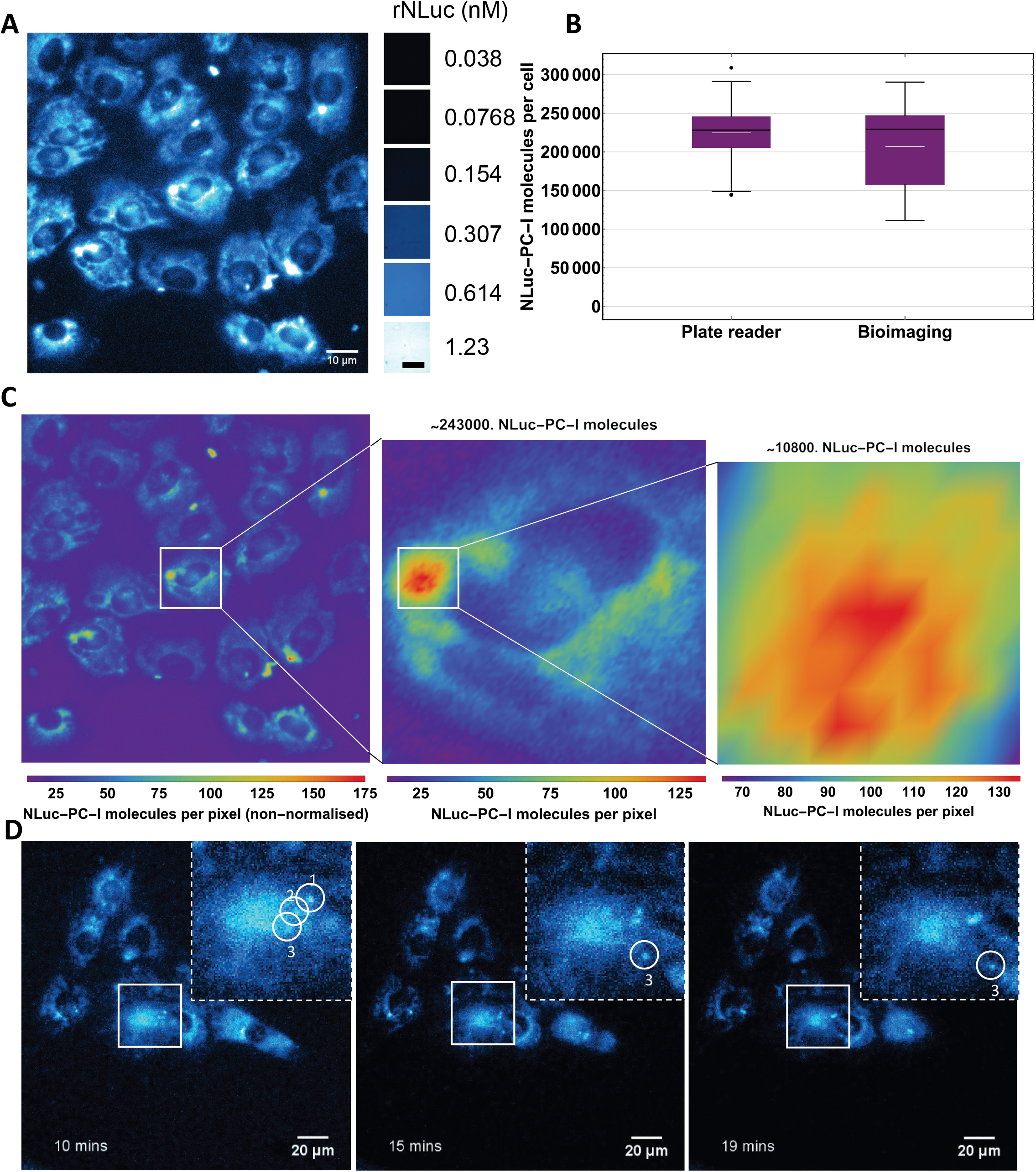
Quantitation at single cell and sub-cellular levels. **A)** Bioluminescence imaging of *nluc::Col1a2* cells. Colourised bioluminescence image of NLuc-PC-I immediately following addition of the substrate Furimazine to 21 cells. Images were taken every minute for 5 mins, and the data summed. Adjacent, summed images of rNLuc used for quantitative correlation, scale bar represents 100 μm. Analysis of images is shown in Supplementary Fig 4. **B)** Quantitation of NLuc-PC-I molecules per cell using bioluminescence imaging of 21 cells were compared to estimates from all plate reader measurements. The black line within the box shows the median value, and the white dash the mean value. The fences show maximum and minimum values (excluding outliers). **C)** Scaled image of A) showing the number of NLuc-PC-I molecules per pixel. NLuc-PC-I was found to be concentrated in puncta, quantitation of a single subcellular vesicle containing 10,800 molecules at a concentration of 0.231 mg/mL or 0.479 μM, assuming a spherical vesicle. **D)** Snapshots of *nluc::Col1a2* cells imaged over time in Supplementary Movie 1. Inserts show magnified images of individual puncta.

Intracellular NLuc-PC-I was also imaged over time at high temporal resolution, allowing for dynamic protein quantitation of NLuc-PC-I in moving vesicles (Fig. 3D and Supplementary Movie 1). It was possible to track the movement of individual vesicles and to estimate their size and number of NLuc molecules they contained. We recorded time-lapse images of the cells (recording for 20 mins at 1-minute intervals, Supplementary Fig. 4A) and noticed that the luminescence from some puncta increased during 20 mins whereas light levels coming from other puncta remained constant and others faded (Supplementary Fig. 4D, E). Furthermore, the intensity of light from puncta was greater than that from the endoplasmic reticulum (ER), and, the intensity of light emanating from the ER decreased during the time series. Presumably these results are explained by NLuc-PC-I exiting ER and being transported to sites within the cell for storage, degradation, or secretion.

### Circadian fluctuations of procollagen-I

We noted that the distribution of NLuc-PC-I luminescence differed from cell to cell (Fig. 3A). It has recently been shown that PC-I levels in tendon fluctuate rhythmically during 24 hours under the control of the circadian clock^6^. Therefore, to explore the possibility that the variation in NLuc levels observed in individual cells could reflect differences in PC-I levels in cells at different stages of the circadian cycle, we synchronised *nluc-Col1a2* cells every 4 hours (Fig. 4A) and measured luminescence as a function of time post-synchronisation. Intracellular NLuc-PC-I exhibited a strong circadian rhythm as shown by MetaCycle^16^ (23.9 hours) and a low Benjamini-Hochberg^17^ q-value (9×10^-10^) (Fig. 4B). The time of peak levels of intracellular luminescence was 12.2 hours post synchronisation (estimated to be circadian time CT0), which aligns well with observations of peak PC-I levels in tendon in vivo^6^. These findings provided direct evidence that the circadian clock influences the synthesis of NLuc-PC-I in *nluc::Col1a2* cells. Rhythmic fluctuations were also observed for secreted NLuc-PC-I, having a period of 27.6 hours (3 s.f.) and a q-value 7×10^-5^ (Fig 4C). Here, overall NLuc levels increased relative to the time after synchronisation, presumably because of PC-I accumulation in the culture medium during the recording period. Fluctuations in NLuc in the matrix fraction were not 24-hour rhythmic (Fig. 4D), this was presumably a result of NLuc-collagen accumulation in the form of fibrils and transport of cleaved N-propeptides.

**Figure 4:**
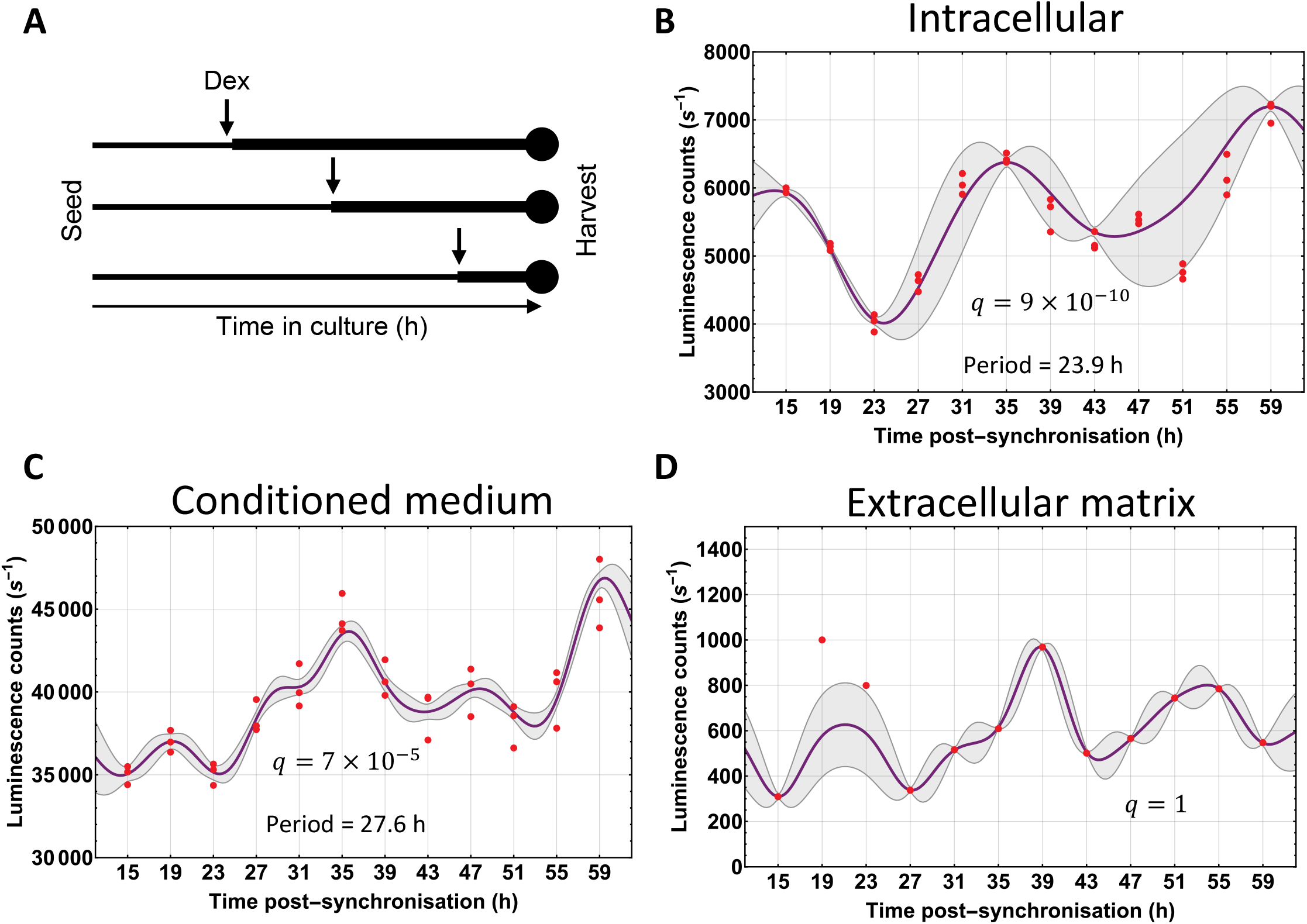
Circadian fluctuations in cellular NLuc-PC-I. **A)** Schematic of experiments performed to assess circadian fluctuations in PC-I in *nluc::Col1a2* cells whilst maintaining consistent cell numbers. **B)** The levels of cellular NLuc-PC-I activity over 48 hours, MetaCycle analysis indicated a 23.9-hour periodic fluctuation in cellular procollagen. Graph shows n=3 independent replicate data points (red), Gaussian process predicted function (purple), and standard deviation of the Gaussian process (grey). **C)** The levels of secreted NLuc-PC-I activity over 48 hours. MetaCycle analysis indicated a 27.8-hour periodic fluctuation in secreted NLuc-PC-I. The cellular and secreted NLuc-PC-I levels follow the same pattern. Graph shows n=3 independent replicates. **D)** The incorporation of NLuc-collagen-I into the extracellular matrix was assessed following decellularisation at each time. Whilst levels fluctuated over 48 hours, MetaCycle analysis did not indicate a periodic incorporation of NLuc-collagen-I into the extracellular matrix. Graph shows n=3 independent replicates.

### NLuc-PC-I response to known collagen modulators

Next, we assessed the ability of *nluc::Col1a2* cells to respond to known modulators of collagen-I. As a first experiment, we showed that blocking protein synthesis with cycloheximide brought about a 90% reduction in levels of NLuc-PC-I in conditioned medium and cells (Supplementary Fig. 5A). The secretory pathway inhibitors Brefeldin A and Monensin both caused inhibition of NLuc-PC-I secretion (Supplementary Fig. 5B). Treatment with Brefeldin A, unlike Monensin, resulted in accumulation of intracellular NLuc-PC-I, which is in line with the fact that Brefeldin A is known to induce a strong ER stress response^18^. Encouraged by these results, we next sought to determine if collagen-DyProQ could be used to evaluate the function of the known anti-fibrotic therapeutics Nintedanib^19^ and Pirfenidone^20^. Using doses which did not significantly impact on cell growth (Supplementary Fig. 6A) we observed a reduction in both secreted and cellular NLuc-PC-I (Supplementary Fig. 5C, D). As a further means of evaluating collagen-DyProQ, the *nluc::Col1a2* cells were treated with the profibrotic growth factors, TGF-β 1, 2, and 3. Treatment with TGF-β 1, and 3 for 72 hours showed strong induction of NLuc-PC-I levels in both cellular and secreted collagen (Supplementary Fig. 5) without significant effect on cell survival (Supplementary Fig. 5). We transfected NIH3T3 cells with a vector expressing NLuc under the control of a Smad-responsive element (Supplementary Fig. 5G) and flow sorted the transfected cells (Supplementary Fig. 6C, D). The selected cells were then treated separately with TGF-β1, 2 and 3 (Supplementary Fig. 5H). We showed that TGF-β2 had a smaller effect on collagen levels compared to TGF-β1 and TGF-β3, which correlated with the degree of SMAD activation by TGFβ ligands.

### High-throughput screen of procollagen modulating compounds

Screens to identify therapeutics that modulate collagens have been performed using hydrophobic dyes that bind multiple collagens^21^, and assays that assess collagen-I transcription^22^ or secretion in overexpression models^23^. However, *nluc::Col1a2* cells preserve endogenous control over collagen transcription, translation, and secretion, with retain responses to circadian cues and TGF-β, and thereby offered new possibilities to identify new collagen-modifying compounds. Using collagen-DyProQ to screen a commercial library of 1,971 off-patent FDA-approved compounds, we identified compounds that increased and decreased PC-I synthesis and secretion (Fig. 5). We measured NLuc-PC-I levels in cells and conditioned medium after 24 and 72 hours, controlled for cell viability (Fig. 5A). Luminescence reads for all compounds are shown (Fig. 5B, Supplementary Fig. 7). Comparisons of secreted NLuc-PC-I after 24 hours relative to both DMSO-treated controls and pre-treatment NLuc-PC-I levels (Fig. 5C) identified 49 compounds that resulted in ≥ 6-fold reduction in secreted NLuc-PC-I and 45 compounds that caused ≥ 2-fold increase. These 94 ‘hits’ were then screened for effects on rNLuc luminescence activity. Of the 49 first-round inhibitors, 9 reduced rNLuc activity (Fig. 5D) and were discarded from further studies. In contrast, the 45 compounds that induced PC-I secretion did not alter rNLuc activity (Supplementary Fig. 8A).

**Figure 5:**
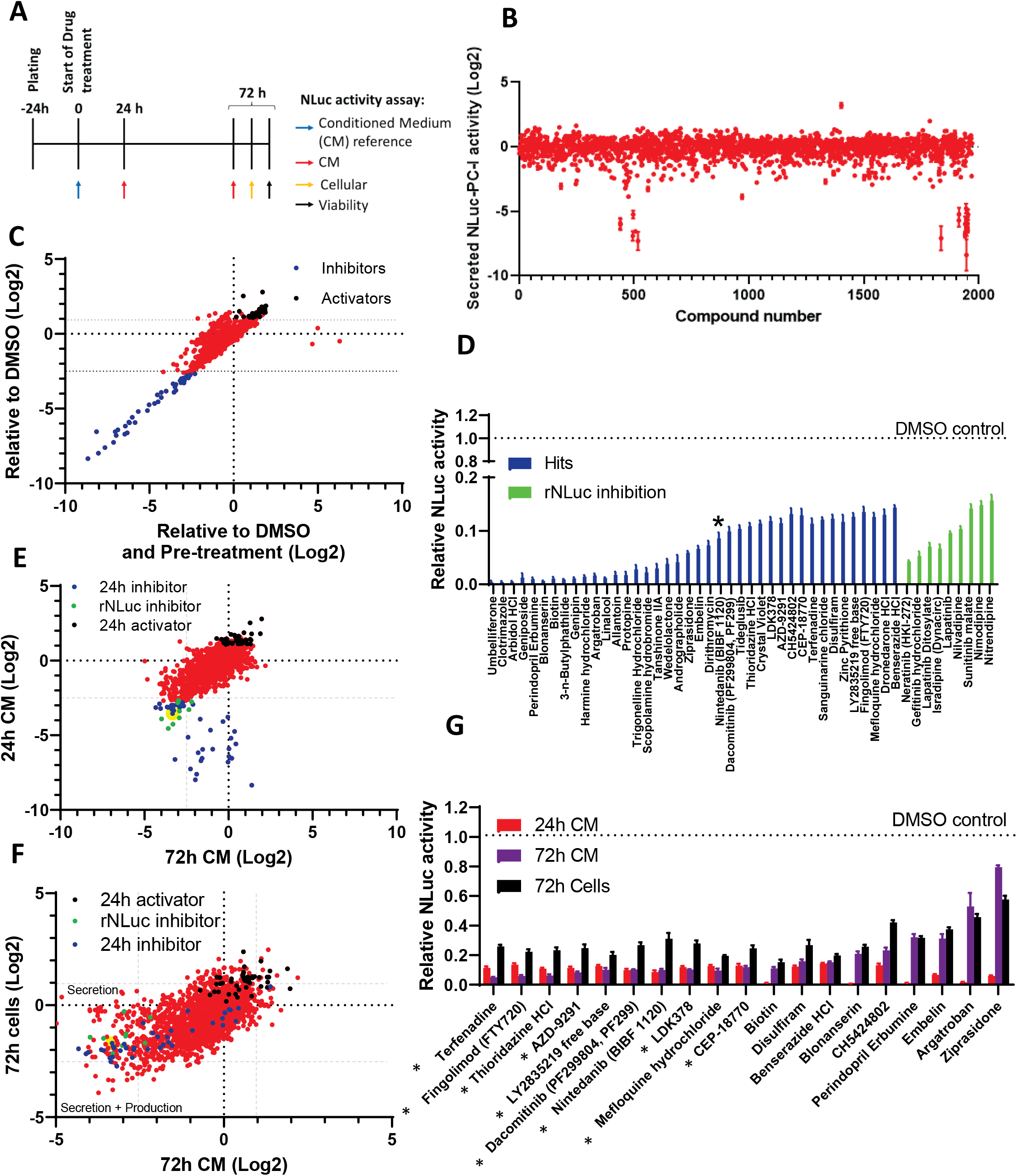
Drug screen for collagen inhibitors/activators. **A)** Experimental set up for drug screen. **B)** The effects of all 1971 compounds on secreted NLuc-PC-I after 24 hours treatment, NLuc-PC-I levels relative to DMSO controls are shown. Error bars show the standard deviation of 4 repeat measurements for each well. **C)** Comparison of two normalisation approaches to identify procollagen modulating therapeutics after 24 hours treatment. Activators were determined by ≥ 2-fold increase in secreted NLuc-PC-I activity. Inhibitors were determined by ≥ 6-fold reduction of secreted collagen. **D)** The 49 inhibitors identified in C) were further screened for their effect on rNLuc activity. Approximately 18% (9/49 compounds) of these hits reduced rNLuc activity. These inhibitors were excluded from further analysis. Asterisk shows the approved anti-fibrotic therapeutic, Nintedanib. Bars show the mean ± SD of n=4 repeat measurements of each treatment. **E)** Comparison of the inhibitors and activators, identified in C) after 24h and 72h incubation showed good correlation between 24h and 72h treated samples. Nintedanib was identified as one of a number of compounds that effectively reduced NLuc-PC-I secretion at both time points and is indicated in yellow. Some hits showed lesser effects after 72h treatment which may reflect compensation to, or, breakdown of, these compounds over time. Compounds that inhibited rNLuc activity showed similar responses at both 24h and 72h in conditioned medium. **F)** Comparison of the effects of compounds on NLuc-PC-I activity within cells and in the conditioned medium, identify additional compounds which equally target both collagen production and secretion, and those that have a greater impact on the secretion of NLuc-PC-I. **G)** The effects of 19 compounds, selected from D, at each time point is shown. Asterisk denotes compound with corresponding dose response data shown in supplementary figure 9. Bars show the mean ± SD of n=4 repeat measurements for each treatment.

Comparison of inhibitory and activating effects after 24 hours and 72 hours treatment identified those compounds with sustained effects (Fig. 5E), and included the approved anti-fibrotic, Nintedanib. However, some of the initial hits at 24 hours were found to lose efficacy over time (Supplementary Fig. 8B), whilst other affecters only influenced NLuc-PC-I levels after prolonged treatment. These compounds were also eliminated from further studies. To gain insight into the mechanisms of how the shortlisted compounds modulate collagen secretion we compared how they affected cellular and secreted levels of NLuc-PC-I (Fig. 5F). We observed a strong linear relationship between cellular and secreted NLuc-PC-I (Supplementary Fig. 7C) i.e. compounds that decreased cellular NLuc-PC-I also tended to decrease levels of secreted NLuc-PC-I, and this trend was observed across the entire screen. We identified 7 compounds that displayed reproducible activation of NLuc-PC-I secretion in culture medium at 24 and 72 post-treatment, and cells at 72 hours post-treatment (Supplementary Fig. 8B).

Some inhibitors had a greater impact on secretion than on cellular NLuc-PC-I levels (including Nintedanib (Fig. 5F)) whilst others reduced collagen secretion without affecting cellular levels, implying inhibition of the collagen secretory pathway. We validated the positive hits from this initial screen by assessing whether they exhibited dose dependent effects on secreted and cellular NLuc-PC-I levels (Fig. 5G). This produced a final list of 10 compounds that showed strong dose-dependent inhibition of PC-I synthesis and secretion (Supplementary Fig. 9).

### Identification of procollagen-modulatory pathways

Hierarchical clustering of effect size at 72 hours from the screen was used to identify pathways that modulate collagen secretion. The heatmap in Fig. 6A illustrates how the lead compounds fell into three categories: i) those that caused up-regulation of NLuc-PC-I levels in cells and culture medium after 72 hours treatment (the top ~1/3^rd^ of compounds shown), ii) those that caused a decrease in NLuc-PC-I levels in cells and culture medium (shown in the bottom half of the heatmap), and those that caused an increase in cellular NLuc-PC-I and a decrease in secreted NLuc-PC-I (shown in the centre of the heatmap as ‘down regulators’ in the culture medium and ‘up regulators’ in the cell).

**Figure 6:**
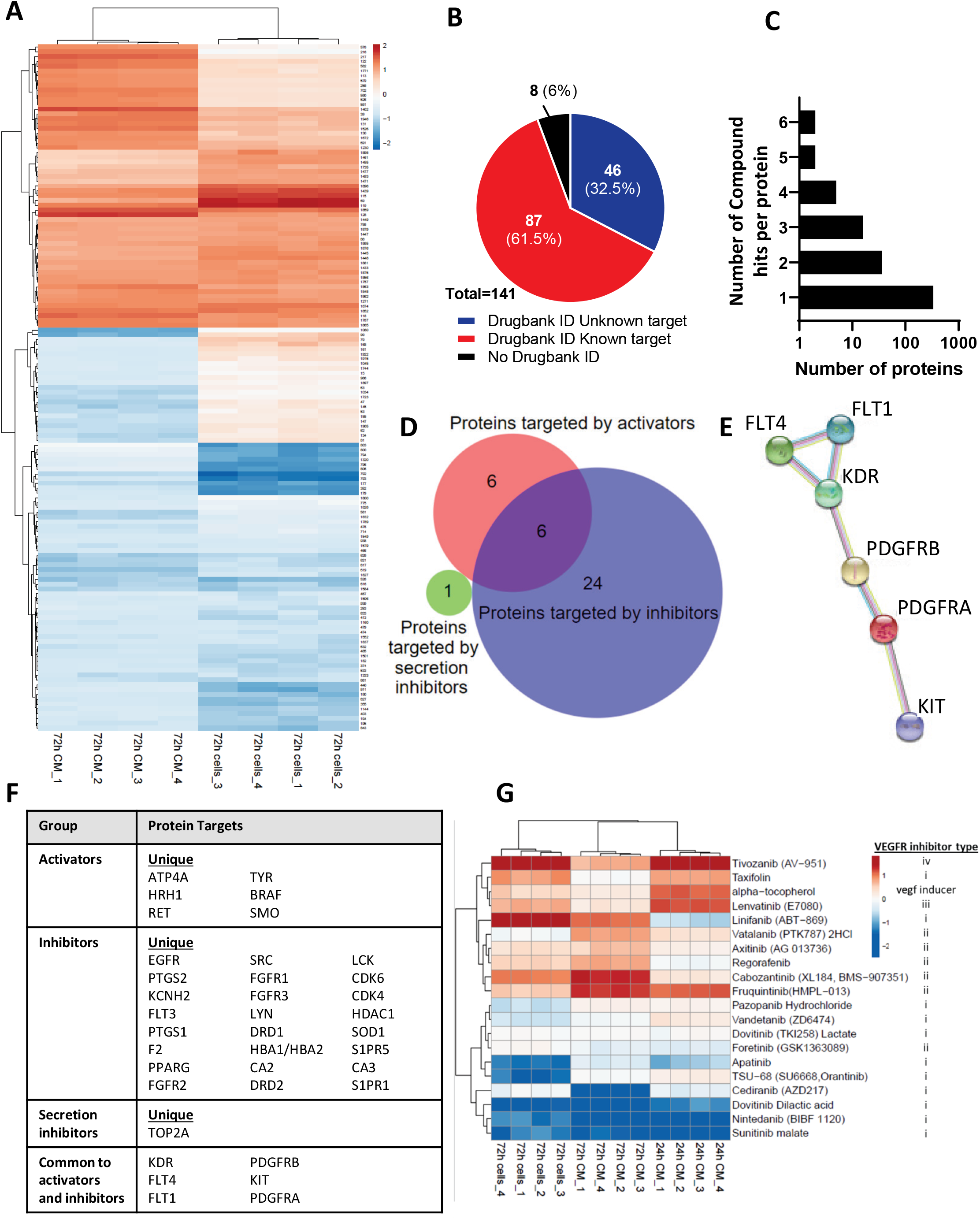
VEGFR inhibitors are common modulators of collagen production. **A)** Heatmap of cellular and secreted NLuc-PC-I levels in inhibitor treated *nluc::Col1a2* cells with ≥ 6-fold reduction or ≥ 2-fold increase in NLuc activity, replicate measurements for each treatment are shown. Three types of collagen modulators were identified; compounds that activate or inhibit collagen production/secretion or those that suppress secretion but not cellular NLuc-PC-I levels. **B)** Using the cut offs in A, 143 compounds identified, there were 141 unique compounds. These were searched against the DrugBank database in order to identify target proteins. **C)** The compounds with known protein targets tended to identify single protein targets, however some targets were hit by multiple compounds. **D)** In order to identify pathways that were targeted whilst altering collagen levels, only protein targets that were targeted by at least two different compounds were included in the analysis. **E)** Of interest were six proteins targeted by both activators and inhibitors of NLuc-PC-I levels. PDGF receptors, VEGF ligand receptors and the stem cell factor (SCF) receptor c-Kit. **F)** Table of common proteins targeted by at least 2 compounds. **G)** Heatmap of all VEGFR targeting compounds, both inhibitors and activators of collagen secretion were identified. Inhibitors of collagen production was aligned with type I inhibitors of VEGFR2.

Of the 143 hits (141 unique compounds), DrugBank identifiers were available for 133, and 87 of these had known protein targets (Fig. 6B). The majority of these proteins were only targeted by single compounds (Fig. 6C). By comparing proteins targeted by at least two compounds in the library screen, we identified 30 proteins that when targeted reduced NLuc-PC-I levels, 12 proteins that when targeted led to elevated NLuc-PC-I levels, and one protein (TOP2A) that was targeted by two compounds (Epirubicin and Idarubicin) and inhibited NLuc-PC-I secretion (Fig. 6D). Surprisingly, there were some proteins that were associated with both reduction and increase in NLuc-PC-I levels (Fig. 6D), depending on the compound in the collection. These proteins included VEGFR1, 2 and 3, PDGFRa and b, and KIT all of which are known to be functional interactors (Fig. 6E and F). We assembled the 20 compounds in the screen that are known to target VEGFR pathways and tabulated the result of the screen (Fig. 6G). This analysis showed that compounds with type I inhibition mechanisms tended to decrease collagen secretion whereas those with type II, III and IV inhibition mechanisms elevated collagen levels.

## Discussion

In our study we developed a method for dynamic protein quantitation (DyProQ) of endogenous proteins. CRISPR-Cas9 mediated insertion of NLuc into the target protein of interest is central to the method. Inserting *nluc* into the gene locus ensures that the normal regulatory elements are maintained. Furthermore, the brightness of NLuc in the presence of Furimazine meant that the use of exogenous expression is not necessary. Therefore, DyProQ will be widely applicable to the study of endogenous protein dynamics. Others have measured protein dynamics using fluorescence correlation spectroscopy^24^ or by using surrogate markers of transcription^25^; however, these lack scalability, and often require exogenous expression of reporters. Using PC-I as a test protein we could, with high precision, determine the number of molecules being synthesised, transported in vesicles, and secreted into the culture medium. Collagen-DyProQ is ~10^5^ – 10^6^ times more sensitive than the conventional chemical method of estimating collagen and has utility across different platforms from bioluminescence microscopy to plate reader-style detection for high-throughput screening. Using this method, we quantified PC-I levels in individual cells and up to 20,000 cells, and we demonstrated the circadian regulation of PC-I synthesis in fibroblasts and the induction of PC-I in the presence of TGF-β, especially TGF-β3. When imaging the concentration of PC-I in individual cells we discovered that cells concentrate PC-I in compartments potentially in preparation for fibril formation. We also screened a library of FDA-approved drugs, whereby we identified off-patent compounds that can be investigated for use in regulating collagen levels in the treatment of fibroproliferative diseases, and we could infer from these data that there are key pathways that can both promote and suppress collagen secretion.

The site of insertion of NLuc into the target protein sequence is likely to have a major bearing on the normal synthesis, trafficking and secretion of the protein of interest. In our study, we chose to place NLuc at the N-terminus of the proα2(I) chain. The trimeric PC-I molecule comprises two proα1(I) and one proα2(I) polypeptide chains; therefore, each NLuc-PC-I molecule carries one NLuc tag. The assembly and subsequent zippering of the trimeric procollagen molecule is initiated by sequences in the C-terminal of each chain^26^. Furthermore, the major triple helical domain of the molecule is particularly sensitive to mutations that change the repeating Gly-X-Y structure, as shown by studies of osteogenesis imperfecta^27^. Therefore, we chose to insert NLuc at the N-terminus of the molecule, and specifically in the proα2(I) chain. The green fluorescent protein has previously been located at this position without interfering with trafficking of the protein and subsequent assembly into fibrils^28^. PC-I is converted to collagen by removal of N- and C-terminal propeptides by procollagen N- and C-proteinases, respectively. Removal of the C-propeptides is required for fibril assembly^29^. However, removal of the N-propeptides is not required for fibril formation^30^ and a proportion of collagen molecules retain N-propeptides in the extracellular matrix^31^. Of particular note, failure to remove the N-propeptides of PC-I results in skin hyperextensibility and joint hypermobility in people with the Ehlers-Danlos syndrome type VII^32^. Therefore, in our study, we chose not to engineer out the N-proteinase cleavage site in the proα2(I) chain so as to maintain the physiological functions of the N-propeptide and to approximate, as near as possible, the normal synthesis, secretion and procollagen-handling behaviour of cells.

The insertion of the GFP11 peptide, a 6 histidine spacer and the NLuc sequences into the N-propeptide of proα2(I) chain was tolerated by PC-I, as shown by i) the presence of PC-I in the culture medium, ii) comparison of PC-I secretion from *nluc::Col1a2* and non-edited cells, and iii) comparison with published values of PC-I secretion (~200,000 procollagen molecules per cell per hour^33^). The high photon output of NLuc combined with bioluminescence microscopy made it possible to count the number of PC-I molecules in transport vesicles in the secretory pathway, and to record the movement of the vesicles by time-lapse by microscopy. We identified vesicles in which the numbers of NLuc-PC-I molecules remained constant during 20 mins, and others in which numbers increased and decreased. These findings provide insights into the possibility that PC-I molecules are delivered to these transport vesicles either for secretion, storage or degradation. This approach also showed that the concentration of PC-I in some transport carriers was 3 orders of magnitude higher than the critical concentration of collagen required for fibril formation^29^, and 5 times higher than the surrounding ER concentration. Therefore, cells concentrate procollagen molecules in preparation for collagen fibril formation. Our ability to measure the number of PC-I molecules in individual cells enabled a time-series study of procollagen synthesis, in which we showed that the synthesis of PC-I was rhythmic with a ~24-hour period, and thereby confirmed previous proteomic data that the synthesis of PC-I is under circadian clock control^6^.

We used collagen-DyProQ to screen a library of off-patent FDA-approved drugs, for which information was available on the proposed protein targets. We identified compounds that increased and decreased NLuc-PC-I secretion. We generated a top 10 list of compounds that decreased procollagen secretion, each with dose-dependent responses. A notable inclusion in this list was Nintedanib, which is a potent small molecule inhibitor of the receptor tyrosine kinases PDGF receptor, FGF receptor and VEGFR^34^ and used in the treatment of pulmonary fibrosis^35^. A proposed mechanism of action is reduced TGF-β stimulated collagen secretion by primary lung fibroblasts^34^. Others included Thioridazine, a first-generation antipsychotic drug and a potential anti-inflammatory^36^; Mefloquinine, used to prevent malaria but has suspected psychotic side effects^37^; CEP-18770 (Delanzomib), a proteasome inhibitor with potential anti-tumour activity^38^; Dacomitinib, an EGFR tyrosine kinase inhibitor being investigated for treatment of non-small-cell lung cancer; LDK378 (Ceritinib), a potent ALK inhibitor used in the treatment of non-small-cell lung cancer^39^; Terfenadine (which produces the metabolite, Fexofenadine) is a second-generation antihistamine that inhibits TNF signalling and is a therapeutic against inflammatory arthritis^40^; Fingolimod is a first-in-class sphingosine-1-phosphate receptor modulator and immunosuppressant that is approved for the treatment of relapsing-remitting multiple sclerosis^41^; AZD-9291 (Osimertinib), is a mutant-selective EGFR inhibitor used to treat non-small-cell lung cancer^42^ and, Benserazide is a peripheral decarboxylase inhibitor that increases the amount of levodopa crossing into the brain and its subsequent conversion to dopamine, and is used in combination with other drugs in the treatment of Parkinson’s disease. We propose that these 10 compounds are candidates for repurposing in the treatment of fibrosis.

In contrast to the PC-I secretion inhibitors that have anti-tumour, anti-fibrotic or anti-inflammatory effects, the 7 compounds that increased NLuc-PC-I secretion (Misoprostol, Levulinic acid, Helicid, Hyodeoxycholic acid, deoxycholic acid and Piperine) have beneficial effects in digestion, pregnancy, cosmetics, and the food industry (e.g. Piperine is a major bioactive ingredient in pepper) and therefore targeted different biochemical pathways. Interestingly, Helicid is used in the preparation of the anti-fibrotic therapeutic, Pirfenidone, which may counteract some of expected effects on collagen secretion and production. Given that collagen I production is under the influence of circadian control mechanisms it was fascinating that Epirubicin and Idarubicin (which targeted NLuc-PC-I secretion but not cellular levels) targeted the DNA topoisomerase, TOP2A. TOP2A has been shown to regulate the period length of Bmal1 transcriptional oscillation^43^. However these effects could also be due to suppression of the TGF-β/Smad pathway^44^.

Some proteins were commonly targeted by both collagen inhibitors and collagen activating compounds (Fig. 6D). These 6 proteins (FLT4, FLT1, KDR, PDGFRB, PDGFRA and KIT) are known to interact. Therefore, therapeutic targeting of this network could be problematic in controlling collagen levels. Whilst there appears to be a trend for type I VEGFR inhibitors to suppress NLuc-PC-I secretion, other classes of VEGFR inhibitors induced procollagen secretion. Seventeen of 20 VEGFR inhibitors altered NLuc-PC-I production suggesting a dynamic role of VEGF receptors in controlling collagen production. Nintedanib is approved for clinical use as it has been shown to supress progression of fibrotic disease^45^, however no therapeutics have been shown to reverse fibrosis. Whilst we demonstrate that Nintedanib suppresses collagen production, the fact that targeting VEGFR activity can have the opposite effect on collagen levels might explain the difficulty encountered in the past in identifying effective treatments to reverse fibroproliferative disease.

Collagen-I-DyProQ has immediate applications in studying the synthesis, trafficking, secretion, and degradation of collagen-I caused by mutations in *Col1a1* and *Col1a2*, such as osteogenesis imperfecta, the Ehlers-Danlos syndromes, and Caffey disease. It also has uses in studying the effects on collagen-I synthesis of mutations in genes associated with collagen synthesis, such as FKBP10 and PLOD2 (Bruck syndrome), and BMP1, CREB3L1, CRTAP, P3H1, PPIB, Serpinh1, and TMEM38B (osteogenesis imperfecta). DyProQ could also be used to study the biosynthesis of other collagens, e.g. collagen-II and collagen-XI in Stickler syndrome, collagen-III and collagen-V in the Ehlers Danlos syndrome, collagen-VI in Ullrich congenital muscular dystrophy and Bethlem myopathy, collagen-VII in epidermolysis bullosa, and collagen-IV in sporadic cerebral small vessel disease^46^ and major common diseases including stroke (reviewed by ^47^). DyProQ has wide-ranging applications in studies of other proteins that are expressed at levels too low to be detected by fluorescent protein tagging of the endogenous protein. Finally, mouse models of DyProQ offer the opportunity for whole animal studies.

## Materials and Methods

### Cell Culture

NIH3T3 mouse embryonic fibroblasts and subsequently CRISPR edited cells were maintained in DMEM (Dulbecco’s Modified Eagle Medium) supplemented with heat-inactivated 10% new-born calf serum, 1% l-glutamine, and 1% Penicillin/Streptomycin. The cells were kept at 37°C in humidified incubators with 5% CO_2_. They were passaged using trypsin.

For 96-well plate reader recordings, cells were seeded into a white plastic plate, in cell culture medium described above. Nano-luciferase substrate was then added as required, at the levels of 0.25 μL per 100 μL medium unless otherwise specified.

### Drug Treatments

For drug treatment tests, 500 cells were plated out per well in a 96-well plate for 24 hours prior to the addition of drugs. For concentrations of drug treatment doses see supplementary table 3. For screening the FDA-approved library of 1971 compounds (APExBIO), 1000 cells per well (96-well plate) were plated 24 hours before treatment, 25 μL of conditioned medium was collected as an assay control before washing wells with PBS. Fresh growth medium (90 μL) was added before adding 10 μL of 0.1 mM drug stocks. These were prepared by thawing the entire library at room temperature for 4 hours, plates were vortexed and spun at 100 x *g* for 30 s. Each compound was diluted 1:100 with fresh DMSO to achieve a 0.1 mM stock.

### Generation of Split GFP Expressing Stable Cells

To allow detection of CRISPR edited cells we included a split GFP tag developed in the Bo Huang lab^48^. The sfGFP1-10 barrel was synthesised and cloned into a lentiviral vector (Vectorbuilder), and further subcloned into a CMV driven vector (pLenti CMV V5-LUC Blast (w567-1) was a gift from Eric Campeau (Addgene plasmid #21474 ; http://n2t.net/addgene:21474 ; RRID:Addgene_21474 ^49^)), briefly FLuc was removed from the vector by digesting with BstXI. sfGFP1-10 was PCR amplified with addition of a signal peptide to target expression to the endoplasmic reticulum (ER), using primers in supplementary table 1, and assembled using a Gibson Assembly master mix (NEB). 5 μg pLV-ERsfGfp1-10 was then transfected into 293T cells, along with 2.5 μg VSVG, 2.5 μg pRSV-Rev and 2.5 μg pMDLg/pRRE, using a 3:1 ratio of PEI:DNA, to generate lentivirus, medium was collected 24-48hours post transfection, filtered through a 0.45 μm filter and added to NIH3T3 cells with 8 μg/mL polybrene. After overnight infection fresh medium was added for 8 hours before selecting for 72 hours in 2.5 μg/mL Blasticidin to generate NIH3T3-ERsfGFP1-10.

### CRISPR Editing

The NanoLuciferase sequence was taken from pNL1.1 vector map (Promega) this and the sfGFP11 sequence^48^ were synthesised as a gBLOCK from IDT (Supplementary Table 2). The 5’ and 3’ homology arms were generated by PCR amplification using a repair template previously used to introduce a Dendra2 tag into the *Col1a2* locus^50^ using primers in supplementary table 1. The NanoLuciferase gBLOCK and homology arms were joined using Gibson assembly master mix (NEB) and transformed into Stbl3 bacteria. Resulting in the generation of an NLuc gfp11 *Col1a2* repair template (Supplementary Table 2)

NIH3T3-ERsfGFP1-10 were used to perform CRISPR editing, 1 μg of repair template was transfected into 200,000 cells using a 3:1 ratio Fugene6:DNA (Promega), after overnight transfection cells were grown in fresh medium for 6 hours, cells were then transfected with a *Col1a2* crRNA (ACTTACATTGGCATGTTGCT AGG), tracrRNA and Cas9 (IDT), as previously described^50^. After overnight transection cells were grown for 72 hours in fresh medium. Cells were sorted based on GFP positivity, and expanded before validating the CRISPR knock-in.

### DNA and RNA validation of CRISPR editing

Knock-in of NanoLuciferase was validated initially using a Nano-Glo assay, and then validated at the DNA level by PCR across the gRNA cut site using primers ValF and ValR (Supplementary table 1). Edited cells were trypsinised, pelleted and lysed using the Hotshot DNA isolation method. To further ensure the knock-in, RNA was isolated from knock-in cells and quantitative PCR was performed from the unedited 3’ end of the *Col1a2* transcript into the NanoLuciferase sequence, PCR products were sequenced using Sanger sequencing. Similarly, primers to the 5’ end of the NanoLuciferase sequence and the unedited *Col1a2* region were used to ensure that the reading frame between NanoLuciferase and col1a2 were maintained.

### In-gel detection of NLuc activity

As a further validation of the *nluc::Col1a2* cell line the molecular weight at which NLuc activity could be detected was determined by 1D gel electrophoresis and in-gel detection of NLuc. *nluc::Col1a2* cells were trypsinised, pelleted at 1000 x g for 5 mins and lysed in 8 M urea, 50 mM Tris pH7.5 supplemented with PMSF and phosphatase inhibitors (Sigma). After centrifugation at 12,000 x g for 5 mins, 50 μg protein was loaded onto a 6% Tris-Glycine gel. Proteins were renatured and assayed according to the Nano-Glo^®^ In-Gel Detection System protocol (Promega). Light produced by NLuc was captured using a Chemidoc MP Imager (Biorad).

### Proteomic validation of CRISPR editing

For validation of NLuc integration into the Col1a2 locus, 1L of culture medium from *nluc::Col1a2* cells was collected over the course of 2 weeks, cells were grown as described above. Aliquots were frozen at −80 °C until use. The conditioned medium was flowed through a 5 ml His-Trap fast-flow (GE life sciences) column at a flow rate of 4 ml/min using an NGC chromatography system (Bio-Rad) with a dedicated sample pump. The column was equilibrated in 20 mM Tris-HCl pH 7.4 with 0.15M NaCl (buffer A). The column was washed by mixing 4% buffer B, which was 20 mM Tris-HCl, 0.15 M NaCl and 500 mM imidazole (Ultrapure). Bound proteins were eluted with a step gradient of 4% to 100% buffer B in reverse flow at a flow rate of 2 ml/min. Eluted proteins were collected in 0.5 mL fractions. Twenty microliters of each fraction as mixed with LDS sample loading buffer (Life technologies) without reducing agents and heated at 95 °C for 5 mins then run on a 6% tris-glycine gels. Following Coomassie Blue staining, bands of interest were excised from the gel and dehydrated using acetonitrile followed by vacuum centrifugation. Dried gel pieces were reduced with 10 mM dithiothreitol and alkylated with 55 mM iodoacetamide. Gel pieces were then washed alternately with 25 mM ammonium bicarbonate followed by acetonitrile. This was repeated, and the gel pieces dried by vacuum centrifugation. Samples were digested with trypsin overnight at 37 °C. Digested samples were analysed by LC-MS/MS using an UltiMate^®^ 3000 Rapid Separation LC (RSLC, Dionex Corporation, Sunnyvale, CA) coupled to an QExactive HF (Thermo Fisher Scientific, Waltham, MA) mass spectrometer. Peptide mixtures were separated using a gradient from 92% A (0.1% FA in water) and 8% B (0.1% FA in acetonitrile) to 33% B, in 44 min at 300 nL min-1, using a 75 mm x 250 μm i.d. 1.7 μM BEH C18, analytical column (Waters). Peptides were selected for fragmentation automatically by data dependant analysis. Mass spectrometry were searched using Mascot (Matrix Science UK), against the Swissprot and Trembl database with taxonomy of Mouse selected as well as a custom database including the sequence of NLuc-tagged Col1a2. Data were validated using Scaffold (Proteome Software, Portland, OR).

### Quantitation of absolute collagen levels

Luminescence activity was recorded from known masses of rNLuc protein in culture medium when treated with Furimazine. The same procedure was carried out for *nluc::Col1a2* cells at differing confluence levels. The results can be seen in Fig. 2A. The rNLuc protein has a mass of 54.254 kDa, and therefore 1 μg contains 1.11 × 10^13^ NLuc molecules, and we can convert from mass to concentration. We then used linear regression to generate equations for the relationship between total rNLuc molecule counts and luminescence counts, and between number of cells and luminescence counts (Fig. 2C, D, E, F). This procedure was repeated for cells in bioluminescence imaging.

### Hydroxyproline Assay

The protocol for the hydroxyproline assay to quantify collagen amounts and correlated to luminescence was as follows. *nluc::Col1a2* cells were trypsinised, washed with PBS, counted and pelleted. NLuc-PC-I activity was assessed in a serial dilution of the cell pellets. Matching numbers of cells were pelleted and frozen at −20 °C for the hydroxyproline quantitation. Hydroxyproline was measured using methods previously described^51^. Briefly, 100 μL 6M HCl was added to the cell pellet and incubated at 100 °C overnight. Samples were cooled to room temperature and spun at 12,000 x *g* for 3 mins to remove residual charcoal. Each sample (50 μL) was mixed with chloramine T (450 μL) and incubated at room temperature for 25 mins. Ehrlich’s reagent (500 μL) was added to each sample and incubated at 65 °C for 10 mins. All samples were compared to hydroxyproline standards treated identically. The absorbance of 100 μL was measured a 96-well plate and absorbance at 558 nm read on a H1 plate reader (Biotek).

### Drug screen analysis

Coelentrazine at a final concentration of 3 μM added per well immediately before reading on a bioluminescence plate reader (Neo2, Biotek) with an integration time of 0.1 s. 88 compounds were tested per plate and each plate contained empty wells, DMSO controls and recombinant NLuc for controls. 25 μL of conditioned medium was assayed per well, and after 72 hours treatment cells were washed and 50 μL fresh medium was added to each well. Coelentrazine was added to each well and plates were read to assess cellular NLuc-PC-I activity, as stated for conditioned medium. Upon completion of all NLuc activity assays, 2.5 μL prestoblue (Thermo Fisher) was added to assess cell viability. To assess NLuc activity, 4 reads per well were taken at each assay point. These readings were normalised to either: pre-treatment NLuc activity assays (24 h samples) or to cell viability (72 h). All wells were normalised to the median NLuc activity across each individual plate and then effect size relative DMSO treated wells was assessed. DrugBank identifiers and known protein targets were taken from DrugBank Version 5.1.5^52^. Hits from the drug screen were searched against DrugBank data based on Chemical Abstracts Service (CAS) number or drug name.

### Bioluminescence Imaging

For bioluminescence imaging of recombinant NLuc and *nluc::Col1a2* cells was performed in black walled μ-Plate 96-well plates (iBidi). All imaging was performed in DMEM containing 10% FBS. For imaging rNLuc, wells containing NIH3T3 cells were used to ensure that the focal point was in the same position as when imaging *nluc::Col1a2* cells. Imaging was performed at 37 °C using a 40x oil objective on a Zeiss LSM880 microscope fitted with a Hamamatsu ImageEM electron multiplying CCD. One-minute integration times were used for all samples.

## Acknowledgements

The research was generously funded by Wellcome in the form of a Wellcome 4-year PhD studentship (210062/Z/17/Z) to BCC, and a Wellcome Senior Investigator Award (110126/Z/15/Z) and Wellcome Centre Award (203128/Z/16/Z) to KEK. The proteomics was performed at the Biological Mass Spectrometry Facility in the Faculty of Biology, Medicine and Health (University of Manchester) with the assistance of Stacey Warwood and Ronan O’Cualain. Flow sorting was performed by with the assistance of Dr. Mike Jackson in the Flow cytometry core facility in the Faculty of Biology, Medicine, and Health (University of Manchester). Bioluminescence imaging was performed In the Bioimaging core facility in Faculty of Biology, Medicine and Health (University of Manchester) with help from Dr. Dave Spiller. Assistance with the TGF-β studies was gratefully received from Dr. Stuart Cain. We thank Prof. Oliver Jensen and Dr. Tom Shearer in the Department of Mathematics (University of Manchester) for advice regarding mathematics.

## Author contributions

AP, BCC and KEK conceived the project. BCC and AP designed and performed experiments and interpreted data. AP supervised the experiments. All authors co-wrote the manuscript.

---

**Supplementary figure 1.**
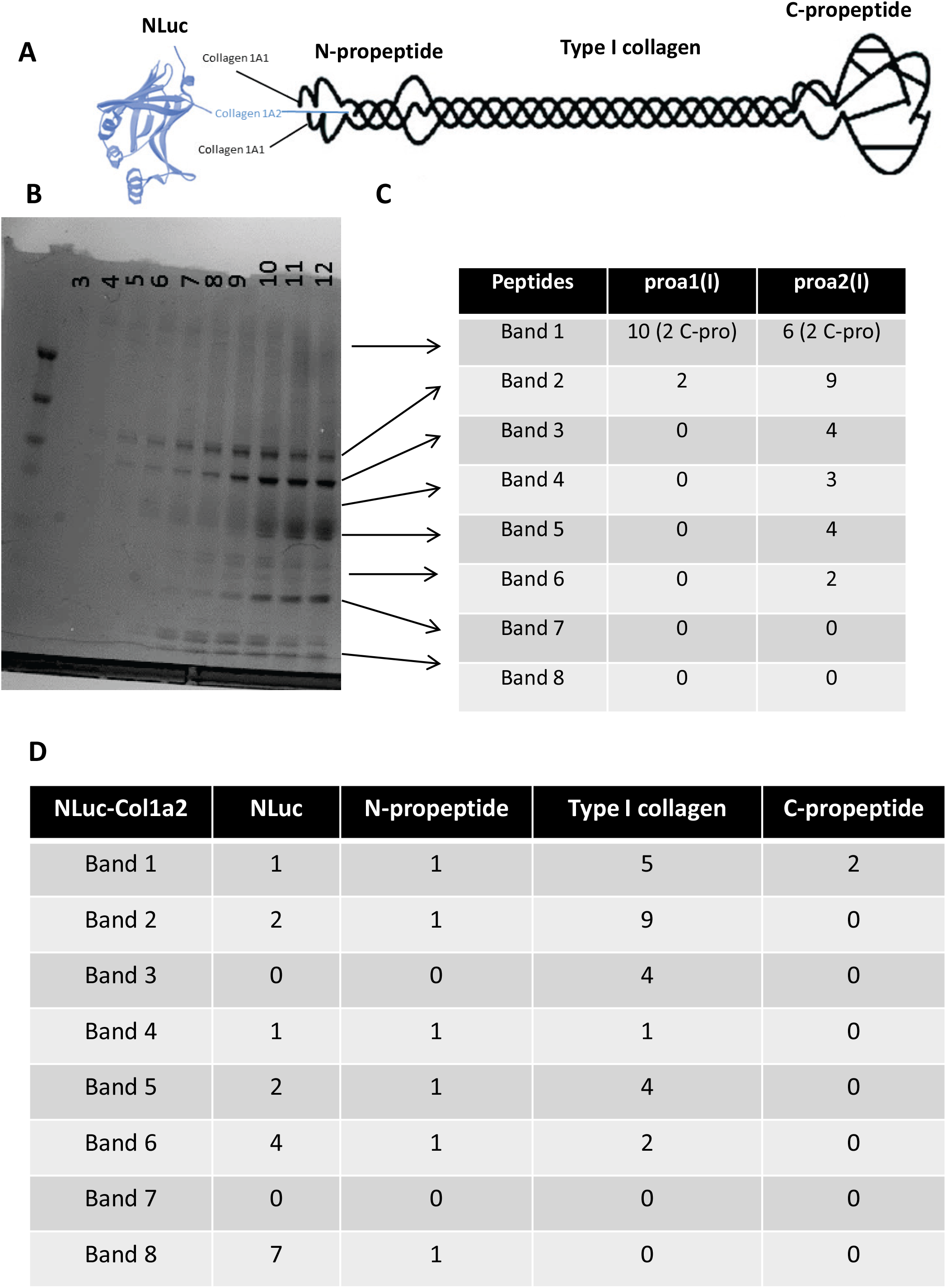
**A)** Diagram of an NLuc tagged type I collagen heterotrimer. **B)** Nickel-affinity purification of NLuc-PC-I from the medium of *nluc::Col1a2* cells, eluted fractions were separated by SDS-PAGE and stained with coomassie blue. **C)** 8 bands were cut from the Coomassie stained gel in B), and proteins identified by mass-spectrometry. Searching against the Swissprot and Trembl database with taxonomy of Mouse selected, the number of peptides identified for Col1a1 and Col1a2 in each of the bands is shown. **D)** The positions of NLuc and Col1a2 peptides are shown and demonstrate that NLuc, N and C propeptides are present at high molecular weight (>250 kDa) indicating the formation of a bona-fide type I heterotrimer. This analysis also identified a peptide spanning NLuc and the N-pro-peptide shown in Figure 1F.

**Supplementary figure 2.**
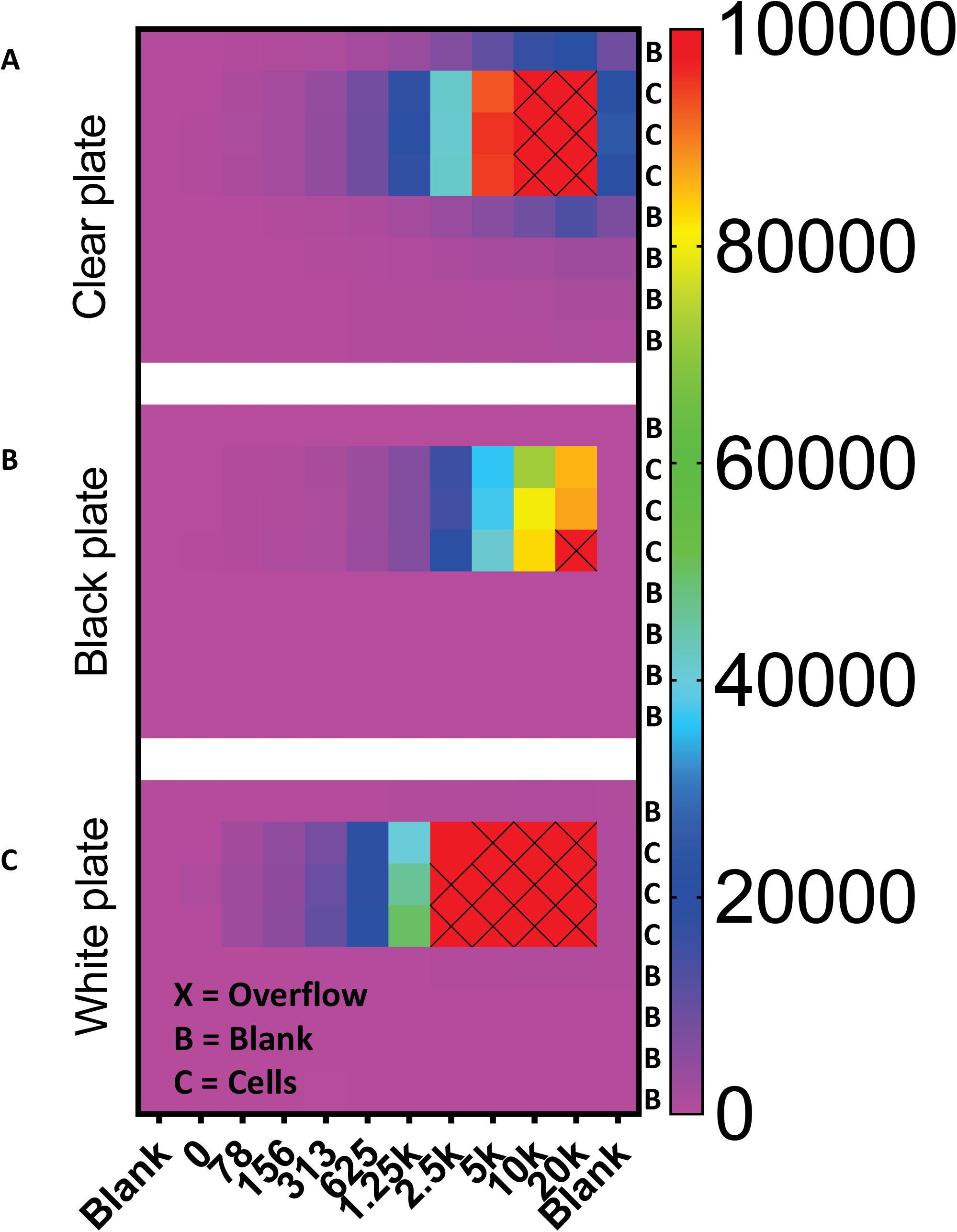
Heatmaps of 96-well plates seeded with different numbers of *nluc::Col1a2* cells. The colour scale (right) indicates the number of counts in each well. Three plates were compared, clear **(A)**, black **(B)** and white **(C)**. Cells were seed in triplicate as indicated. **A)** In clear plates the spill over of luminescence could be observed with large numbers of cells, indicated by significant luminescence being recorded in empty wells neighbouring wells containing cells. **B)** Black plates limited spill over of signal between wells but also suppressed luminescence in the wells. **C)** White plates limited spill over but also enhanced sensitivity of detection.

**Supplementary figure 3.**
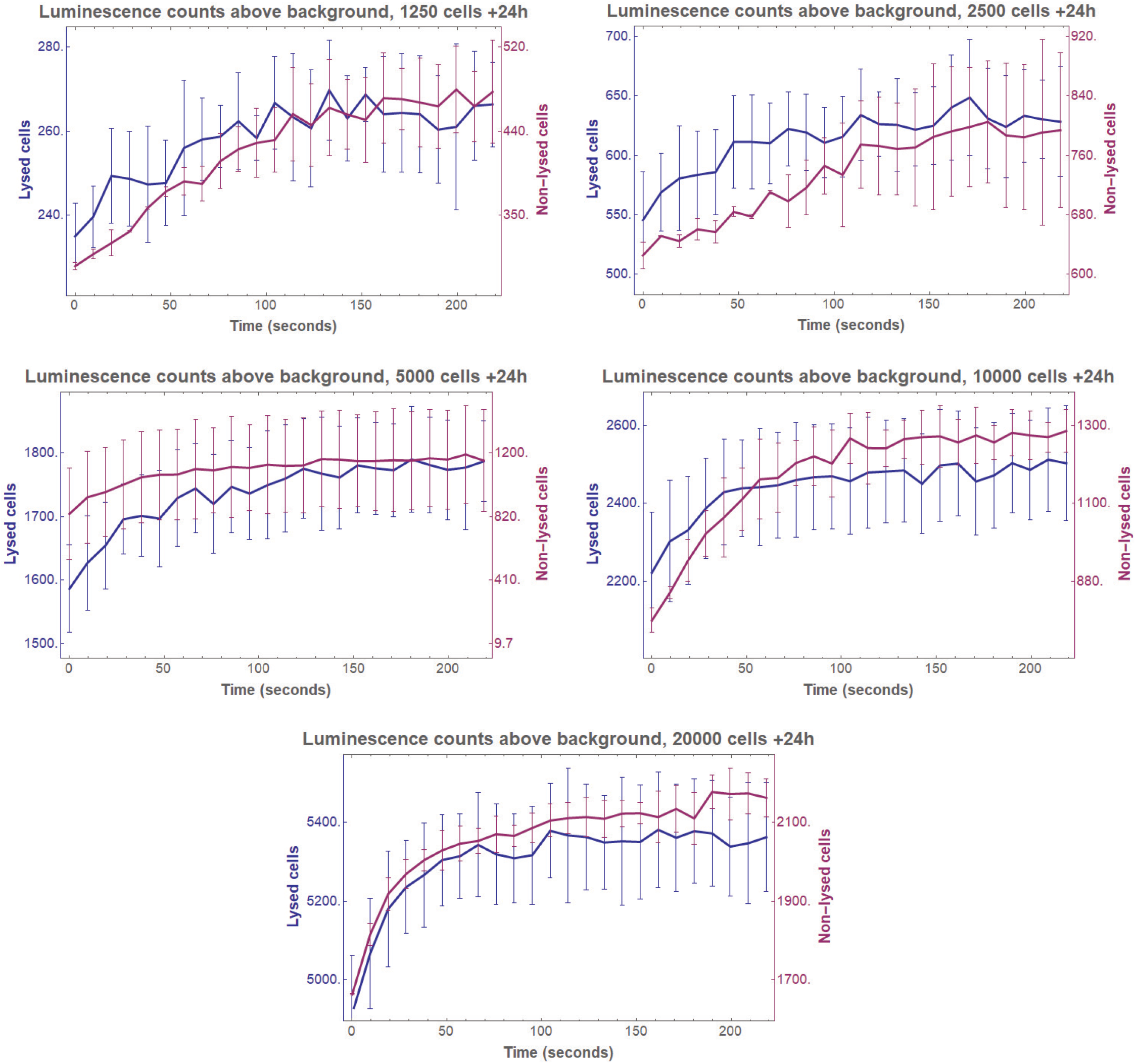
Luminescence counts recorded every 10 second following addition of Furimazine for 1250, 2500, 5000, 10000 and 20000 cells, either in fresh DMEM culture medium (Non-lysed Cells) or after lysis in Nano-Glo buffer (lysed cells). Whilst absolute counts per seconds varied between lysed and non-lysed cells, the time taken to achieve a consistent reading did not alter. The line traces the mean luminescence reading and error bars show the standard deviation of n=4 repeat measurements.

**Supplementary figure 4.**
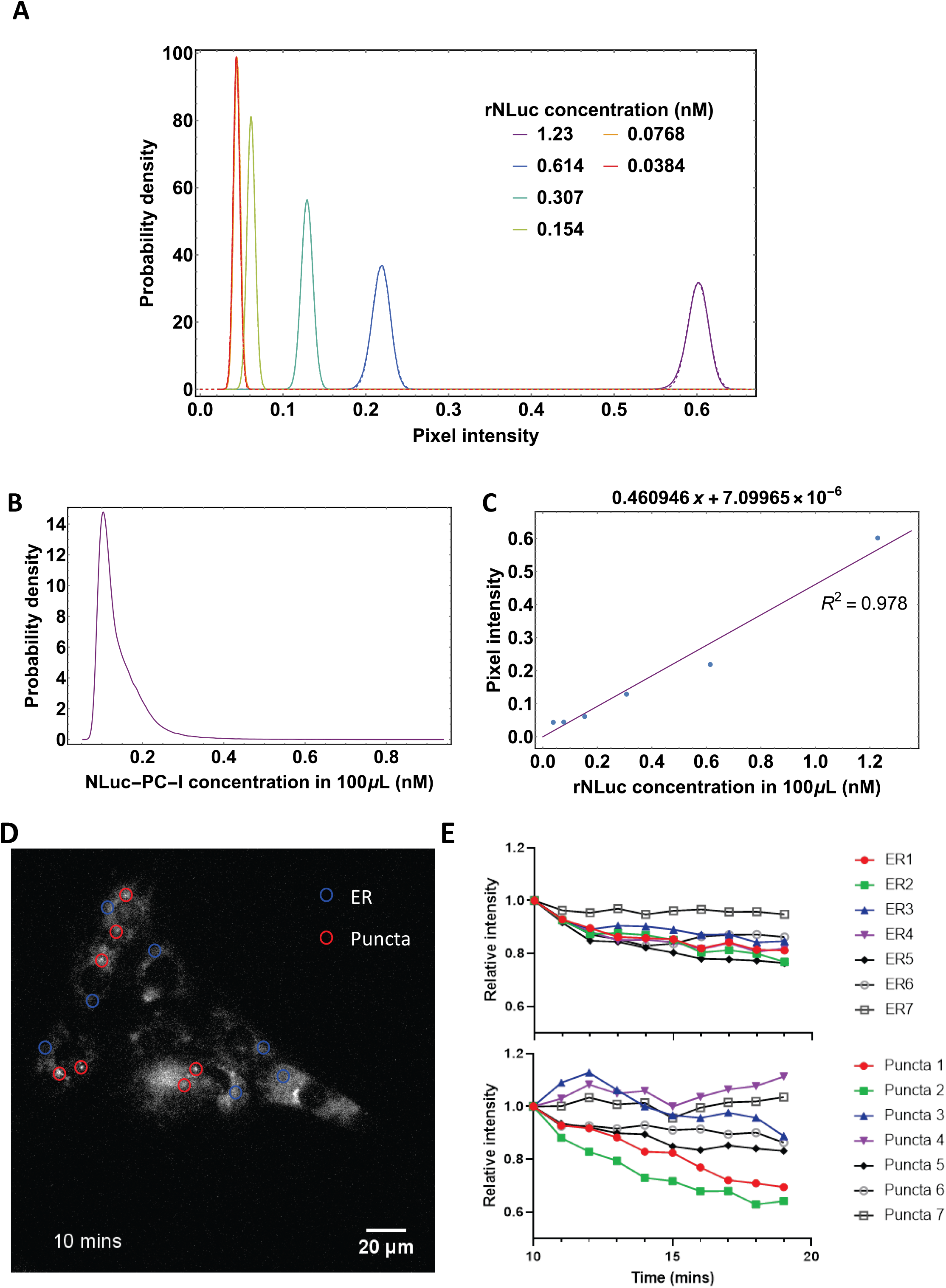
**A)** Probability distribution function of rNLuc pixel intensities at different concentrations (solid), with fitted Gaussian distributions (dashed). **B)** Linear regression of mean pixel intensity from the fitted Gaussian distribution in A) against rNLuc concentration. **C)** Probability density function of pixel intensity of imaged cells in Figure 3A. **D)** The same image as in Fig. 3C with regions of endoplasmic reticulum (ER, Blue circles) and puncta indicated (Red circles). **E)** The change in bioluminescence over 10 mins of imaging in the seven indicated regions of ER and puncta shown in (D).

**Supplementary figure 5.**
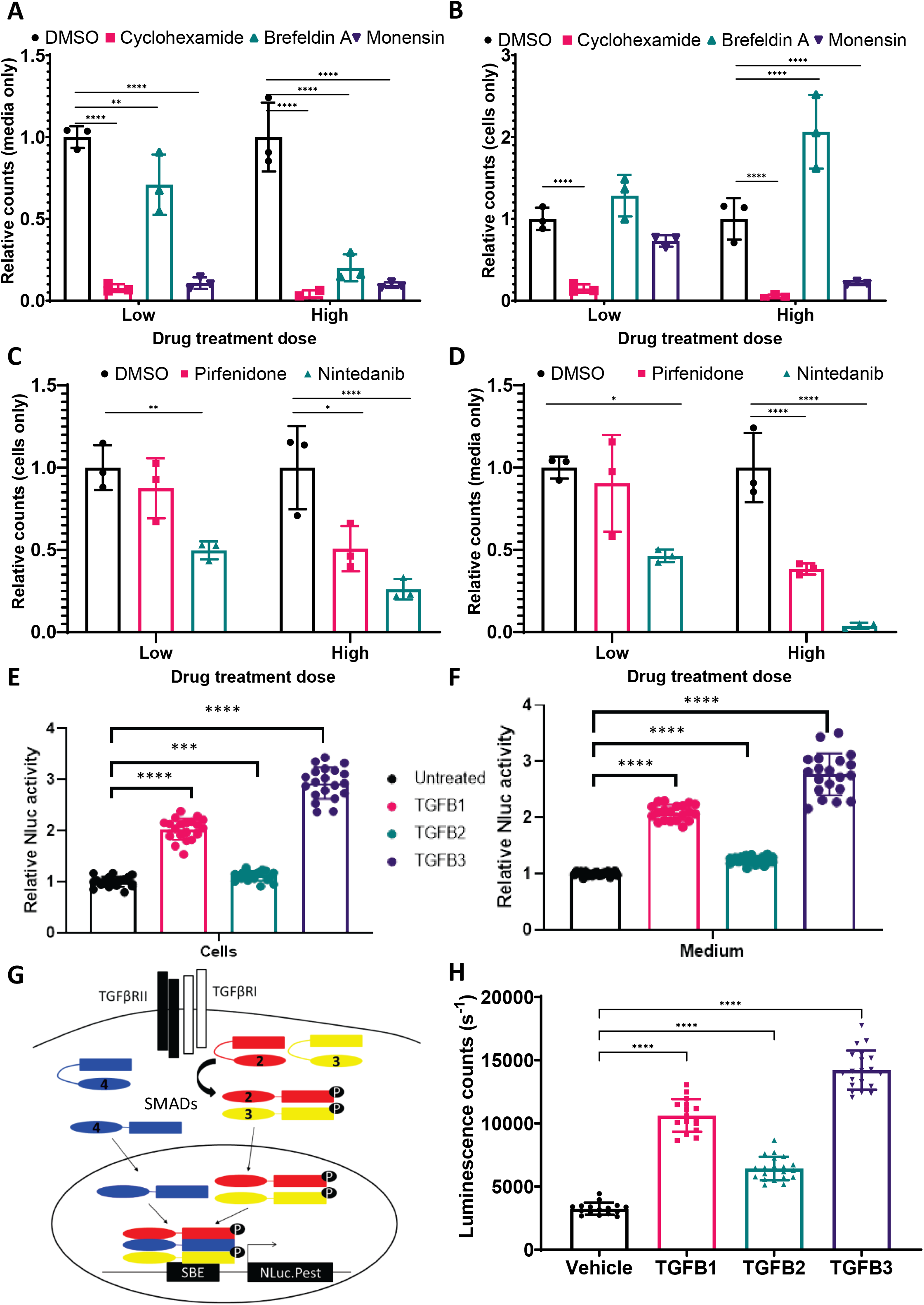
**A)** Cellular luminescence in *nluc::Col1a2* cells treated with cycloheximide or the secretion inhibitors Brefeldin A and Monensin after 24 hours, the doses used are shown in Supplementary Table 3. Bars show the mean ± SD for n=3 independent replicate measurements. **B)** The levels of secreted NLuc after 24 hours treatment are shown, demonstrating each treatment results in inhibition of NLuc-PC-I secretion. Bars show the mean ± SD for n=3 independent replicate measurements. **C)** The effects of the FDA approved therapeutics Nintedanib and Pirfenidone on cellular NLuc activity after 24 hours treatment. Bars show the mean ± SD for n=3 independent replicate measurements. **D)** The effects of Nintedanib and Pirfenidone on NLuc-PC-I secretion with 24 hours treatment. Bars show the mean ± SD for n=3 independent replicate measurements. For charts A-D, * denotes p<0.05, ** denotes p<0.01, *** denotes p<0.001 and **** denotes p<0.0001 paired Student’s t-Test. **E)** Effect of 72 hour treatment of *nluc::Col1a2* cells with TGF-β 1, 2, and 3 treatments on cellular NLuc activity. n=5 independent experiments each with 4 technical repeats. **** indicates p=0.0001 Students paired t-Test, *** indicates p=0.0005 Students paired t-Test. **F)** The effects of TGF-β on secreted NLuc activity. **G)** Schematic of SMAD binding element reporter, SMAD2/3 phosphorylation and activation of SMAD4 following binding of TGF-β ligands to the receptor results in recruitment of SMADs to the SBE which drives NLuc-PEST expression. Following flow sorting of stable lentivirus infected cell lines was detected by flow cytometry, Supplementary Fig 7. **H)** NIH3T3 stably expressing the SMAD binding element driven NLuc reporter, SBE-NLuc-Pest-RFP, demonstrating robust activation of SMADs after 1-hour treatment with TGFB1 and TGFB3, a smaller but significant induction of the SMAD reporter was observed with TGFB2 treatment. n=5 independent experiments each with 4 technical repeats. **** denotes p=0.0001, paired Student’s t-Test.

**Supplementary figure 6.**
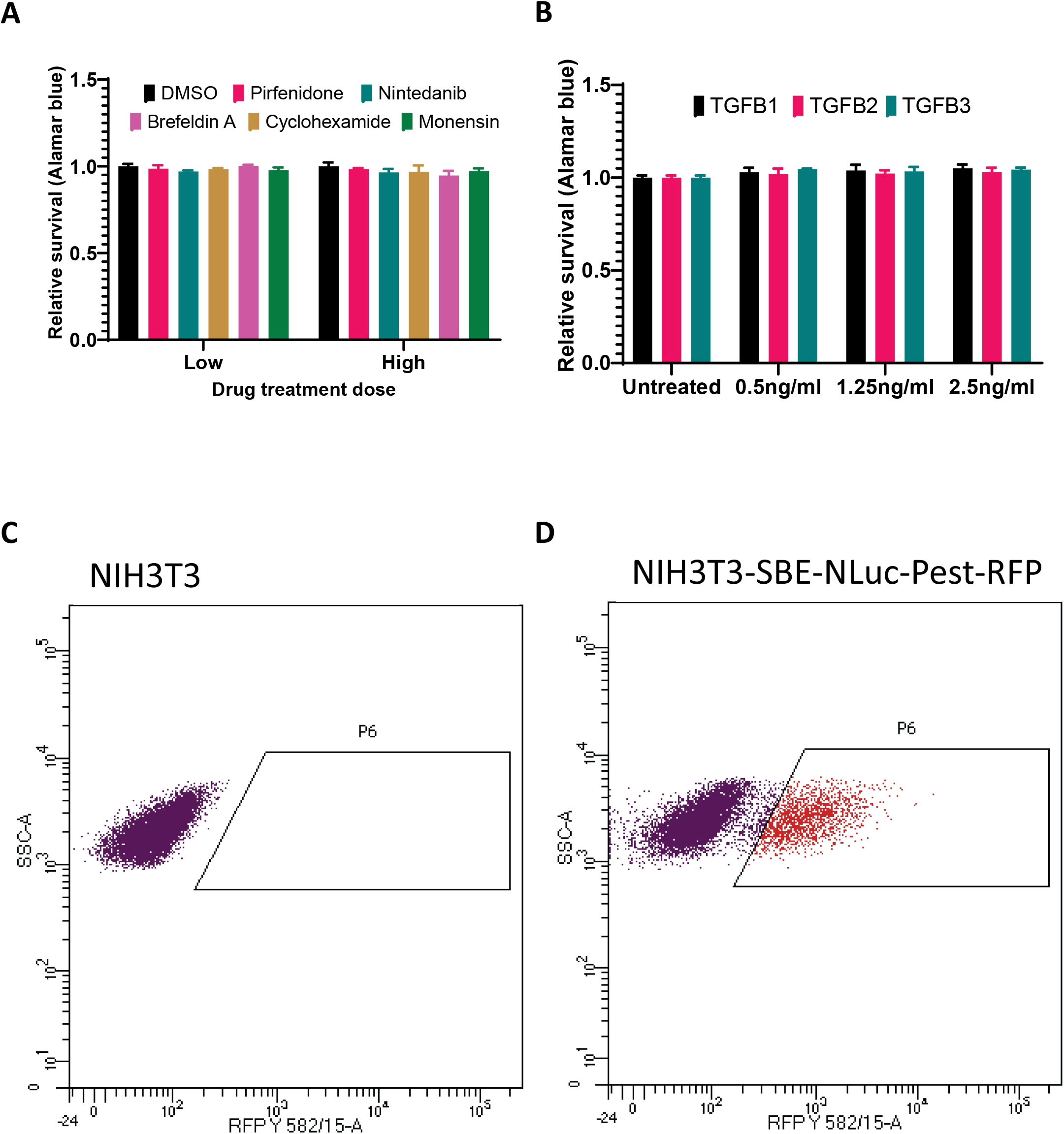
Alamar blue viability assay relative to DMSO treatment for NLuc-PC-I cells treated with **A)** compounds used in Supplementary Fig 6. and **B)** TGF-β 1,2, and 3 after 24 hours. Bars show the mean ± SD, n=3 independent experiments. **C)** Flow sorting of control NIH3T3 cells and **D)** NIH3T3 transduced with the SMAD reporter, SBE-NLuc-Pest-RFP lentivirus. Cells were gated on RFP expression and sorted to generate a stable cell line NIH3T3-SBE-NLuc-Pest-RFP used in Supplementary Fig. 5H.

**Supplementary figure 7.**
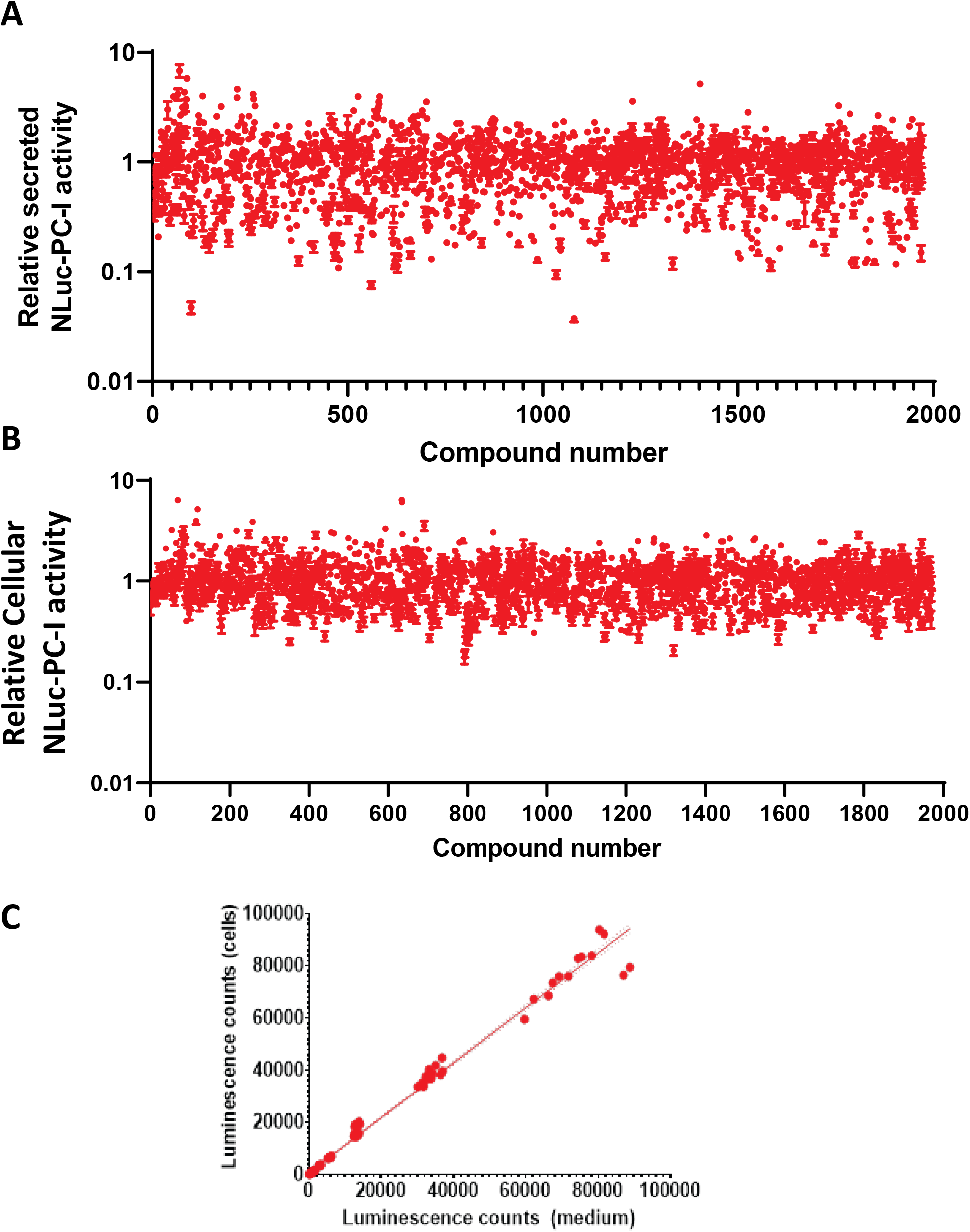
**A)** The effects of all 1971 compounds on secreted NLuc-PC-I after 72 hours treatment, NLuc-PC-I levels are shown relative to DMSO controls, the mean ± SD of 4 repeat measurements for each well are shown. Each effect size was also adjusted for effects on cell viability. **B)** The effects of all 1971 compounds on cellular NLuc-PC-I after 24 hours treatment, NLuc-PC-I levels relative to DMSO controls are shown. Error bars show the standard deviation of 4 repeat measurements for each well. Each effect size was also adjusted to effects on cell viability. **C)** Correlation of cellular and secreted NLuc-PC-I activity using different numbers of cells.

**Supplementary figure 8.**
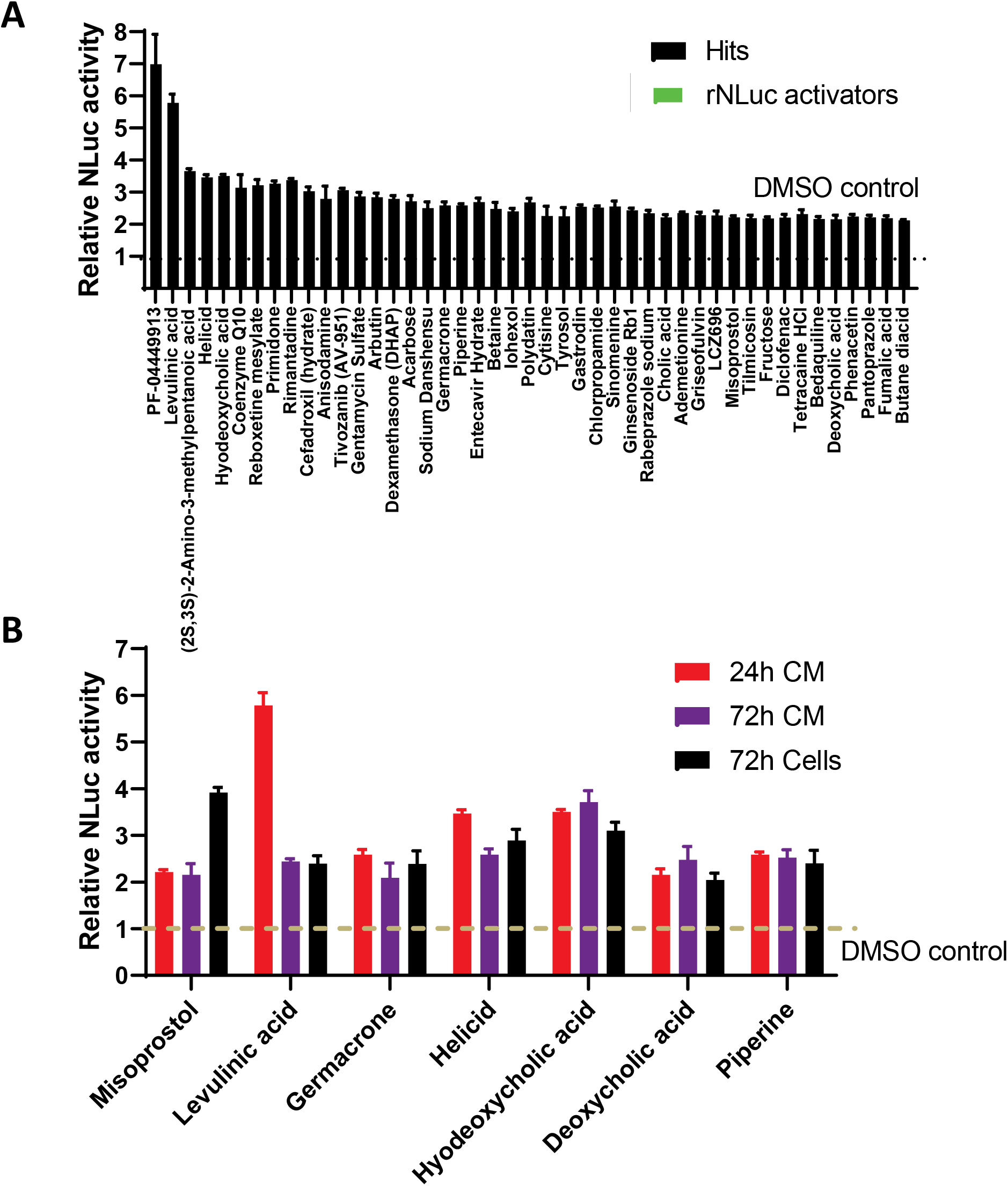
**A)** Plot of the effects of 45 compounds found to induce collagen secretion compared to DMSO after 24 hours treatment in the screen shown in Fig 5B. None of the 45 compounds were found to alter rNLuc activity. Bars show the mean ± SD of 4 repeat measurements for each compound. B) Plot of 7 compounds from A that demonstrated enhanced collagen secretion, compared to DMSO, at 24- and 72-hours treatment and also enhanced cellular procollagen levels. Bars show the mean ± SD of 4 repeat measurements for each compound.

**Supplementary figure 9.**
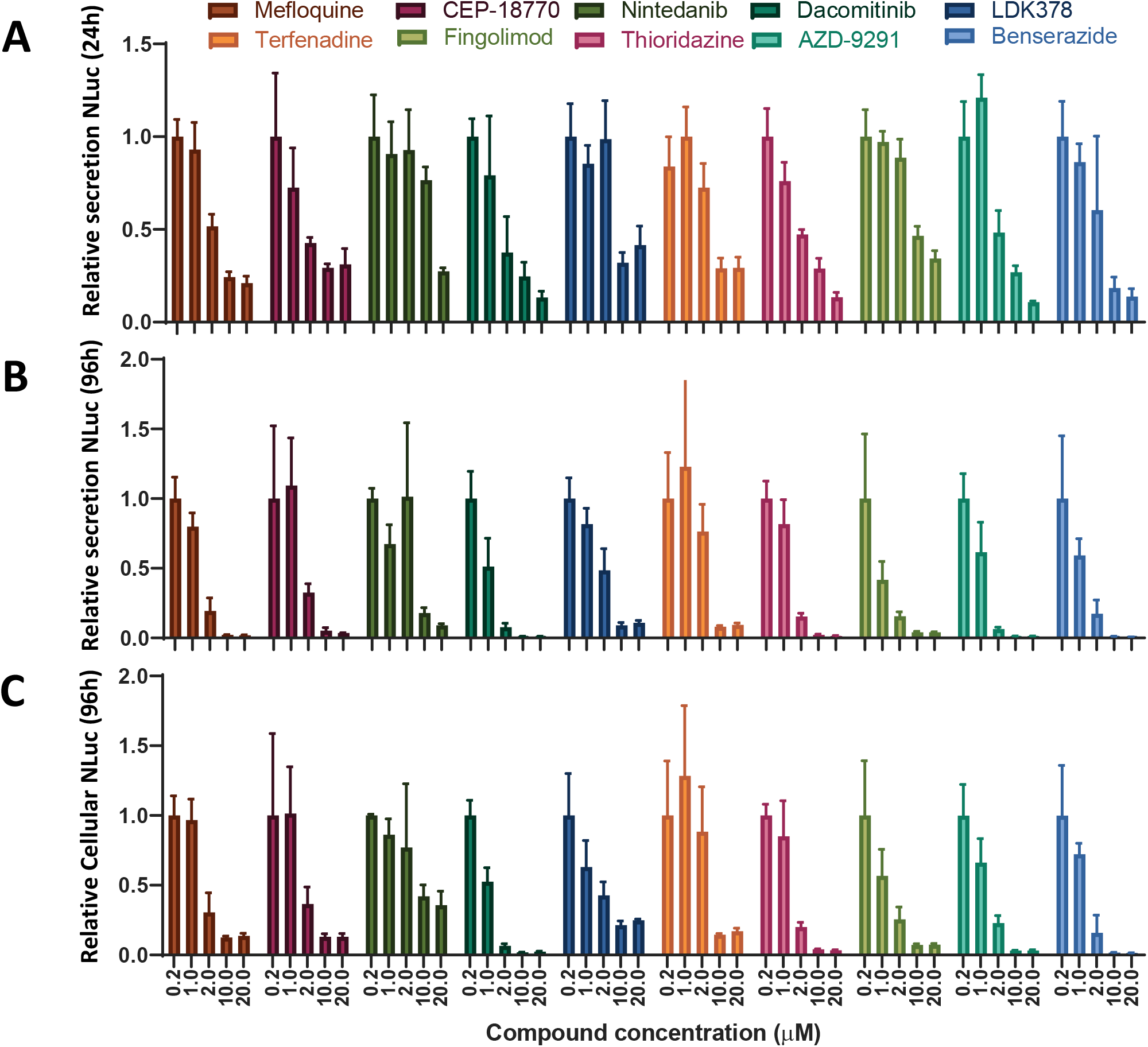
**A)** Dose dependent effects of 10 compounds selected as the most efficient inhibitors of NLuc-PC-I secretion. The levels of secreted NLuc-PC-I after 24 hours treatment, effects were normalised to the lowest dose tested. Bars represent the mean ± SD for n=3 independent repeats each with n=4 repeat measurements for each concentration. **B)** The same wells/cells were assessed for NLuc-PC-I secretion after 72 hours treatment and also cellular NLuc-PC-I levels **(C)**. Bars represent the mean ± SD for n=3 independent repeats each with n=4 repeat measurements for each concentration.

## Supplementary Material

**Supplementary Table 1.**
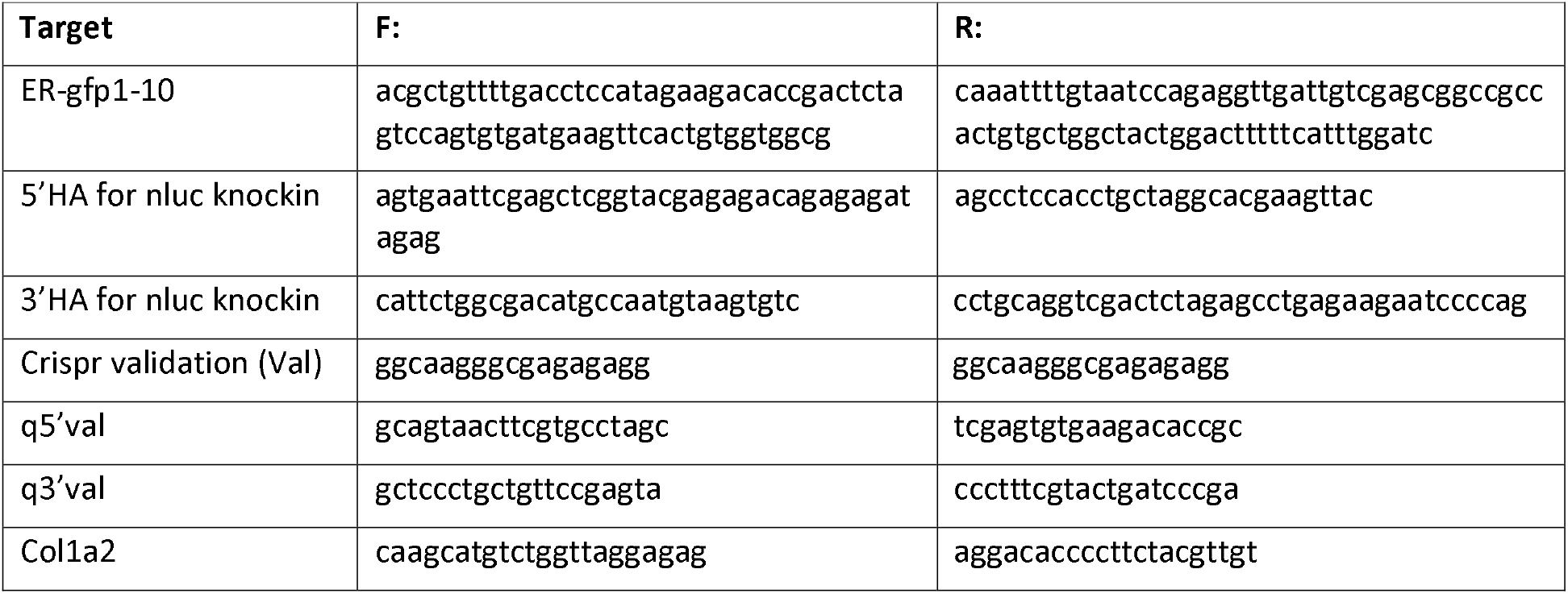
Primers used

**Supplementary Table 2.**
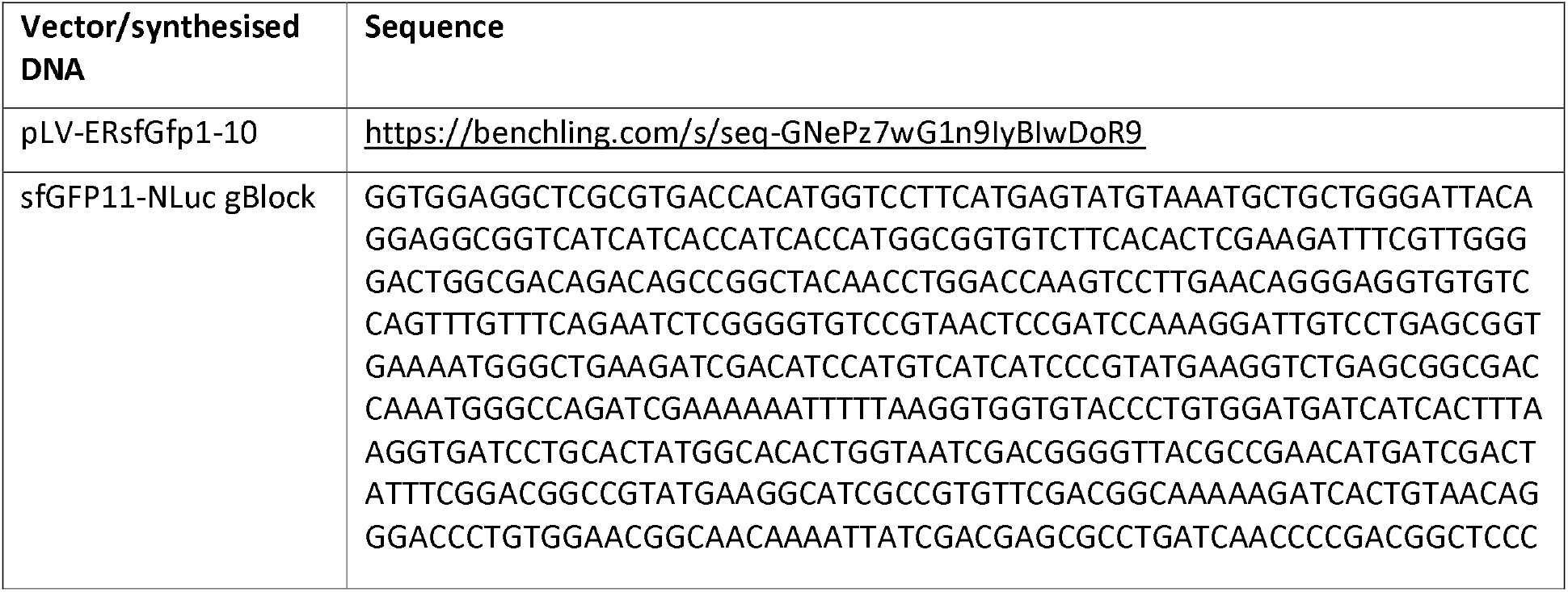
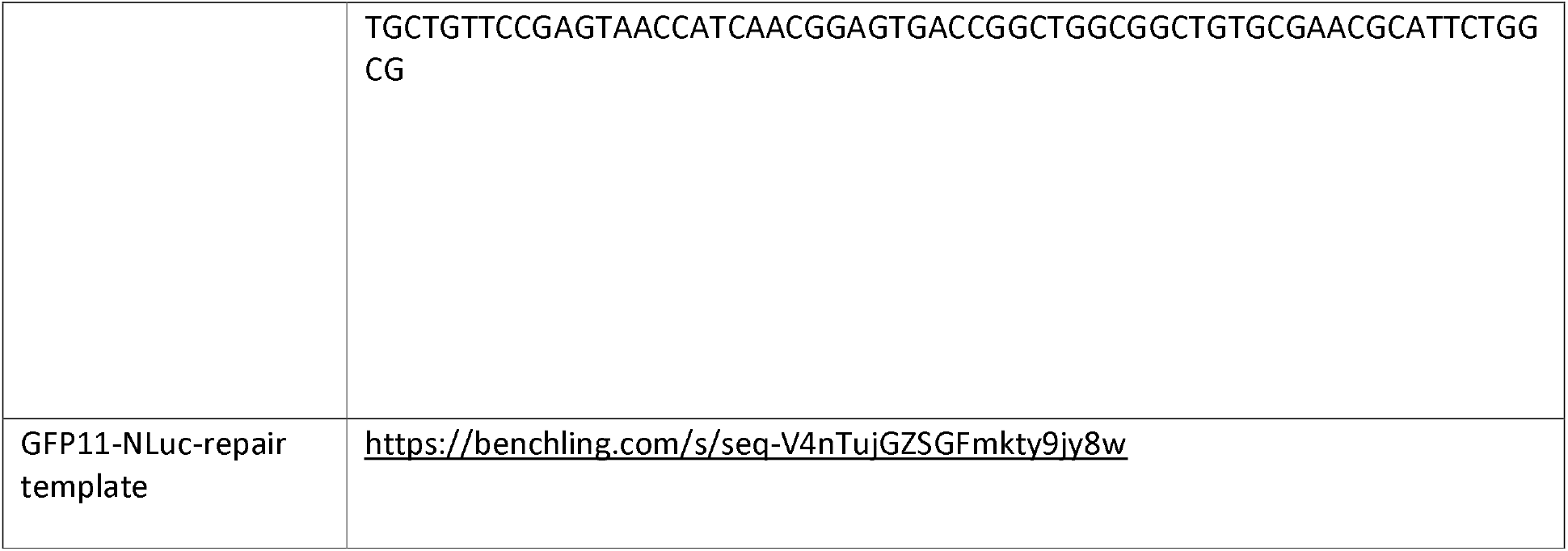
Sequences

**Supplementary Table 3.**
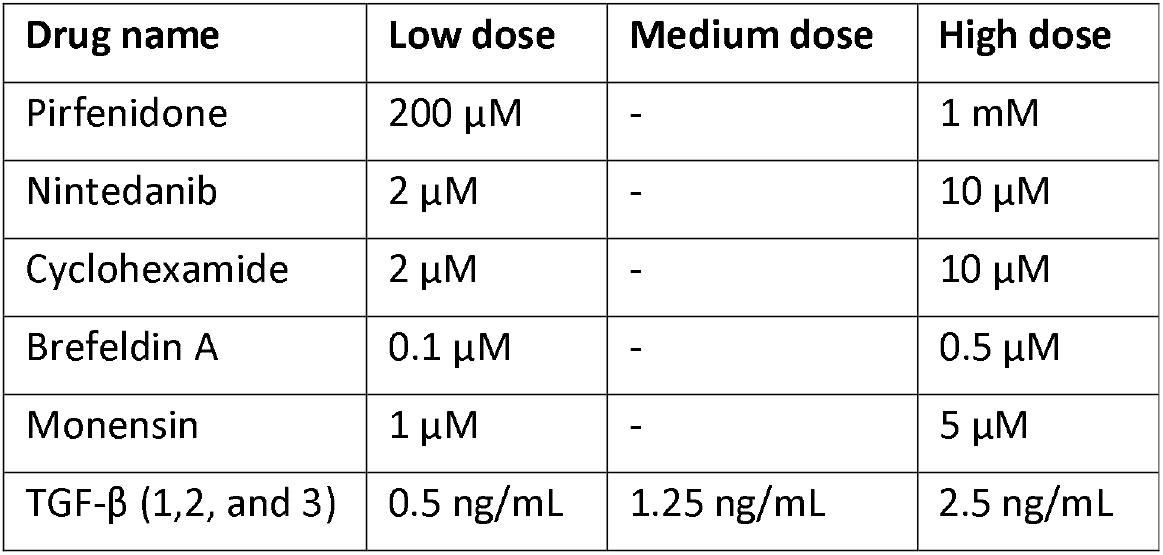
Drug dosage regimes

## References

1 Hall, M. P. et al. Engineered luciferase reporter from a deep sea shrimp utilizing a novel imidazopyrazinone substrate. ACS Chem Biol 7, 1848–1857, doi:10.1021/cb3002478 (2012).

2 Zdzieblik, D., Oesser, S., Baumstark, M. W., Gollhofer, A. & Konig, D. Collagen peptide supplementation in combination with resistance training improves body composition and increases muscle strength in elderly sarcopenic men: a randomised controlled trial. Br J Nutr 114, 1237–1245, doi:10.1017/S0007114515002810 (2015).

3 Bella, J. & Hulmes, D. J. Fibrillar Collagens. Subcell Biochem 82, 457–490, doi:10.1007/978-3-319-49674-0_14 (2017).

4 Kalson, N. S. et al. A structure-based extracellular matrix expansion mechanism of fibrous tissue growth. Elife 4, doi:10.7554/eLife.05958 (2015).

5 Heinemeier, K. M., Schjerling, P., Heinemeier, J., Magnusson, S. P. & Kjaer, M. Lack of tissue renewal in human adult Achilles tendon is revealed by nuclear bomb (14)C. FASEB J 27, 2074–2079, doi:10.1096/fj.12-225599 (2013).

6 Chang, J. et al. Circadian control of the secretory pathway maintains collagen homeostasis. Nat Cell Biol 22, 74–86, doi:10.1038/s41556-019-0441-z (2020).

7 Xu, S. et al. The role of collagen in cancer: from bench to bedside. J Transl Med 17, 309, doi:10.1186/s12967-019-2058-1 (2019).

8 Fang, M., Yuan, J., Peng, C. & Li, Y. Collagen as a double-edged sword in tumor progression. Tumour Biol 35, 2871–2882, doi:10.1007/s13277-013-1511-7 (2014).

9 Wynn, T. A. Common and unique mechanisms regulate fibrosis in various fibroproliferative diseases. J Clin Invest 117, 524–529, doi:10.1172/JCI31487 (2007).

10 Prockop, D. J. & Udenfriend, S. A specific method for the analysis of hydroxyproline in tissues and urine. Anal Biochem 1, 228–239, doi:10.1016/0003-2697(60)90050-6 (1960).

11 Kadler, K. E., Baldock, C., Bella, J. & Boot-Handford, R. P. Collagens at a glance. J Cell Sci 120, 1955–1958, doi:10.1242/jcs.03453 (2007).

12 Brodsky, B. & Persikov, A. V. Molecular structure of the collagen triple helix. Adv Protein Chem 70, 301–339, doi:10.1016/S0065-3233(05)70009-7 (2005).

13 Bentley, J. P. & Hanson, A. N. The hydroxyproline of elastin. Biochim Biophys Acta 175, 339–344, doi:10.1016/0005-2795(69)90011-7 (1969).

14 Canty, E. G. et al. Coalignment of plasma membrane channels and protrusions (fibripositors) specifies the parallelism of tendon. J Cell Biol 165, 553–563, doi:10.1083/jcb.200312071 (2004).

15 Tanzawa, K., Berger, J. & Prockop, D. J. Type I procollagen N-proteinase from whole chick embryos. Cleavage of a homotrimer of pro-alpha 1(I) chains and the requirement for procollagen with a triple-helical conformation. J Biol Chem 260, 1120–1126 (1985).

16 Wu, G., Anafi, R. C., Hughes, M. E., Kornacker, K. & Hogenesch, J. B. MetaCycle: an integrated R package to evaluate periodicity in large scale data. Bioinformatics 32, 3351–3353, doi:10.1093/bioinformatics/btw405 (2016).

17 Benjamini, Y. & Hochberg, Y. Controlling the false discovery rate; a practical and powerful approach to multiple testing. Journal of the Royal Statistical Society. Series B (Methodology) 57, 589–300 (1995).

18 Pickard, A. et al. Preservation of circadian rhythms by the protein folding chaperone, BiP. FASEB J 33, 7479–7489, doi:10.1096/fj.201802366RR (2019).

19 Chaudhary, N. I. et al. Inhibition of PDGF, VEGF and FGF signalling attenuates fibrosis. Eur Respir J 29, 976–985, doi:10.1183/09031936.00152106 (2007).

20 Salazar-Montes, A., Ruiz-Corro, L., Lopez-Reyes, A., Castrejon-Gomez, E. & Armendariz-Borunda, J. Potent antioxidant role of pirfenidone in experimental cirrhosis. Eur J Pharmacol 595, 69–77, doi:10.1016/j.ejphar.2008.06.110 (2008).

21 Tullberg-Reinert, H. & Jundt, G. In situ measurement of collagen synthesis by human bone cells with a sirius red-based colorimetric microassay: effects of transforming growth factor beta2 and ascorbic acid 2-phosphate. Histochem Cell Biol 112, 271–276, doi:10.1007/s004180050447 (1999).

22 Bagchi, R. A., Mozolevska, V., Abrenica, B. & Czubryt, M. P. Development of a high throughput luciferase reporter gene system for screening activators and repressors of human collagen Ialpha2 gene expression. Can J Physiol Pharmacol 93, 887–892, doi:10.1139/cjpp-2014-0521 (2015).

23 Wong, M. Y. et al. A High-Throughput Assay for Collagen Secretion Suggests an Unanticipated Role for Hsp90 in Collagen Production. Biochemistry 57, 2814–2827, doi:10.1021/acs.biochem.8b00378 (2018).

24 Veerapathiran, S. & Wohland, T. Fluorescence techniques in developmental biology. J Biosci 43, 541–553 (2018).

25 Suter, D. M. et al. Mammalian genes are transcribed with widely different bursting kinetics. Science 332, 472–474, doi:10.1126/science.1198817 (2011).

26 Lees, J. F., Tasab, M. & Bulleid, N. J. Identification of the molecular recognition sequence which determines the type-specific assembly of procollagen. EMBO J 16, 908–916, doi:10.1093/emboj/16.5.908 (1997).

27 Marini, J. C. et al. Osteogenesis imperfecta. Nat Rev Dis Primers 3, 17052, doi:10.1038/nrdp.2017.52 (2017).

28 Morris, J. L. et al. Live imaging of collagen deposition during skin development and repair in a collagen I - GFP fusion transgenic zebrafish line. Dev Biol 441, 4–11, doi:10.1016/j.ydbio.2018.06.001 (2018).

29 Kadler, K. E., Hojima, Y. & Prockop, D. J. Assembly of collagen fibrils de novo by cleavage of the type I pC-collagen with procollagen C-proteinase. Assay of critical concentration demonstrates that collagen self-assembly is a classical example of an entropy-driven process. J Biol Chem 262, 15696–15701 (1987).

30 Hulmes, D. J. et al. Pleomorphism in type I collagen fibrils produced by persistence of the procollagen N-propeptide. J Mol Biol 210, 337–345, doi:10.1016/0022-2836(89)90335-5 (1989).

31 Fleischmajer, R. et al. Ultrastructural identification of extension aminopropeptides of type I and III collagens in human skin. Proc Natl Acad Sci U S A 78, 7360–7364, doi:10.1073/pnas.78.12.7360 (1981).

32 Van Damme, T. et al. Expanding the clinical and mutational spectrum of the Ehlers-Danlos syndrome, dermatosparaxis type. Genet Med 18, 882–891, doi:10.1038/gim.2015.188 (2016).

33 Tolstoshev, P. et al. Procollagen production and procollagen messenger RNA levels and activity in human lung fibroblasts during periods of rapid and stationary growth. J Biol Chem 256, 3135–3140 (1981).

34 Wollin, L. et al. Mode of action of nintedanib in the treatment of idiopathic pulmonary fibrosis. Eur Respir J 45, 1434–1445, doi:10.1183/09031936.00174914 (2015).

35 Richeldi, L. et al. Efficacy and safety of nintedanib in idiopathic pulmonary fibrosis. N Engl J Med 370, 2071–2082, doi:10.1056/NEJMoa1402584 (2014).

36 Baig, M. S. et al. Repurposing Thioridazine (TDZ) as an anti-inflammatory agent. Sci Rep 8, 12471, doi:10.1038/s41598-018-30763-5 (2018).

37 Croft, A. & Garner, P. Mefloquine to prevent malaria: a systematic review of trials. BMJ 315, 1412–1416, doi:10.1136/bmj.315.7120.1412 (1997).

38 Wang, M. et al. Delanzomib, a novel proteasome inhibitor, sensitizes breast cancer cells to doxorubicin-induced apoptosis. Thorac Cancer 10, 918–929, doi:10.1111/1759-7714.13030 (2019).

39 Li, S. et al. Ceritinib (LDK378): a potent alternative to crizotinib for ALK-rearranged non-small-cell lung cancer. Clin Lung Cancer 16, 86–91, doi:10.1016/j.cllc.2014.09.011 (2015).

40 Liu, R. et al. Fexofenadine inhibits TNF signaling through targeting to cytosolic phospholipase A2 and is therapeutic against inflammatory arthritis. Ann Rheum Dis 78, 1524–1535, doi:10.1136/annrheumdis-2019-215543 (2019).

41 Wienemann, T. et al. Cryptococcal meningoencephalitis in an IgG2-deficient patient with multiple sclerosis on fingolimod therapy for more than five years - case report. BMC Neurol 20, 158, doi:10.1186/s12883-020-01741-0 (2020).

42 Assadollahi, V., Rashidieh, B., Alasvand, M., Abdolahi, A. & Lopez, J. A. Interaction and molecular dynamics simulation study of Osimertinib (AstraZeneca 9291) anticancer drug with the EGFR kinase domain in native protein and mutated L844V and C797S. J Cell Biochem 120, 13046–13055, doi:10.1002/jcb.28575 (2019).

43 Ogawa, Y., Kawano, Y., Yamazaki, Y. & Onishi, Y. Shikonin shortens the circadian period: possible involvement of Top2 inhibition. Biochem Biophys Res Commun 443, 339–343, doi:10.1016/j.bbrc.2013.11.116 (2014).

44 Gao, Y. et al. TOP2A Promotes Tumorigenesis of High-grade Serous Ovarian Cancer by Regulating the TGF-beta/Smad Pathway. J Cancer 11, 4181–4192, doi:10.7150/jca.42736 (2020).

45 Kolb, M. et al. Nintedanib in patients with idiopathic pulmonary fibrosis and preserved lung volume. Thorax 72, 340–346, doi:10.1136/thoraxjnl-2016-208710 (2017).

46 Rannikmae, K. et al. Common variation in COL4A1/COL4A2 is associated with sporadic cerebral small vessel disease. Neurology 84, 918–926, doi:10.1212/WNL.0000000000001309 (2015).

47 Gatseva, A., Sin, Y. Y., Brezzo, G. & Van Agtmael, T. Basement membrane collagens and disease mechanisms. Essays Biochem 63, 297–312, doi:10.1042/EBC20180071 (2019).

48 Kamiyama, D. et al. Versatile protein tagging in cells with split fluorescent protein. Nat Commun 7, 11046, doi:10.1038/ncomms11046 (2016).

49 Campeau, E. et al. A versatile viral system for expression and depletion of proteins in mammalian cells. PLoS One 4, e6529, doi:10.1371/journal.pone.0006529 (2009).

50 Pickard, A. et al. Collagen assembly and turnover imaged with a CRISPR-Cas9 engineered Dendra2 tag. bioRxiv, doi:10.1101/331496 (2018).

51 Reddy, G. K. & Enwemeka, C. S. A simplified method for the analysis of hydroxyproline in biological tissues. Clin Biochem 29, 225–229, doi:10.1016/0009-9120(96)00003-6 (1996).

52 Wishart, D. S. et al. DrugBank 5.0: a major update to the DrugBank database for 2018. Nucleic Acids Res 46, D1074–D1082, doi:10.1093/nar/gkx1037 (2018).

